# scRNA-seq identifies unique macrophage population in murine model of ozone induced asthma exacerbation

**DOI:** 10.1101/2024.07.23.604740

**Authors:** Jess L. Ray, Joshua Walum, Daria Jelic, Ryelie Barnes, Ian D. Bentley, Rodney D. Britt, Joshua A. Englert, Megan N. Ballinger

**Affiliations:** Division of Pulmonary, Critical Care, and Sleep Medicine, Department of Internal Medicine, Dorothy M. Davis Heart and Lung Research Institute, The Ohio State University, Columbus, OH; Center for Perinatal Research, The Abigail Wexner Research Institute at Nationwide Children’s Hospital, Columbus, OH; Department of Pediatrics, The Ohio State University, Columbus, OH

**Keywords:** Asthma, ozone, macrophage, monocyte, single cell RNA sequencing

## Abstract

Ozone (O_3_) inhalation triggers asthmatic airway hyperresponsiveness (AHR), but the mechanisms by which this occurs are unknown. Previously, we developed a murine model of dust mite, ragweed, and *aspergillus* (DRA)-induced allergic lung inflammation followed by O_3_ exposure for mechanistic investigation. The present study used single cell RNA-sequencing for unbiased profiling of immune cells within the lungs of mice exposed to DRA, O_3_, or DRA+O_3_, to identify the components of the immune cell niche that contribute to AHR. Alveolar macrophages (AMs) had the greatest number of differentially expressed genes following DRA+O_3_, most of which were unique to the 2-hit exposure. Following DRA+O_3_, AMs activated transcriptional pathways related to cholesterol biosynthesis, degradation of the extracellular matrix, endosomal TLR processing, and various cytokine signals. We also identified AM and monocyte subset populations that were unique to the DRA+O_3_ group. These unique AMs activated gene pathways related to inflammation, sphingolipid metabolism, and bronchial constriction. The unique monocyte population had a gene signature that suggested phospholipase activation and increased degradation of the extracellular matrix. Flow cytometry analysis of BAL immune cells showed recruited monocyte-derived AMs after DRA and DRA+O_3_, but not after O_3_ exposure alone. O_3_ alone increased BAL neutrophils but this response was attenuated in DRA+O_3_ mice. DRA-induced changes in the airspace immune cell profile were reflected in elevated BAL cytokine/chemokine levels following DRA+O_3_ compared to O_3_ alone. The present work highlights the role of monocytes and AMs in the response to O_3_ and suggests that the presence of distinct subpopulations following allergic inflammation may contribute to O_3_-induced AHR.

## Introduction

Clinically, asthma is characterized by airway remodeling and episodic exacerbations that lead to increased shortness of breath and wheezing. Asthma exacerbations occur in response to respiratory infections or environmental exposures and result in significant morbidity and even mortality (1). One common environmental exposure that leads to asthma exacerbations is ambient (ground-level) ozone (O_3_). Ground-level O_3_ is an air pollutant formed as a byproduct from the photochemical reactions between primary air pollutants such as volatile organic compounds and nitrogen oxides with UV light (2). Epidemiologic studies have linked exposure to ambient O_3_ to increased use of asthma rescue medications and increased hospital visits (3). Observational studies suggest that asthmatics have increased susceptibility to the adverse effects of O_3_ compared to non-asthmatics (4). However, the cellular and molecular mechanisms by which O_3_ exposure leads to asthma exacerbations are not known.

Many of the clinical manifestations of O_3_ induced asthma exacerbations are due to bronchoconstriction and airway hyperresponsiveness (AHR). AHR is a complex process mediated by increased contraction (or impaired relaxation) of airway smooth muscle (5). Airway smooth muscle hypercontractility is mediated, in part, by inflammatory cells recruited to asthmatic airways. Previous studies have identified recruited granulocytes such as eosinophils, neutrophils, and mast cells as potential mediators of asthmatic AHR in response to O_3_ (6). It has also been observed that there is an increase in other immune cells including alveolar macrophages (AMs) in the airways of asthmatic patients following experimental O_3_ exposure (7). AMs, along with recruited granulocytes, create an inflamed niche around the airways that mediate O_3_ induced AHR.

Current treatment for asthma exacerbations, regardless of the precipitant, includes the use of corticosteroids and bronchodilators. This “one size fits all” approach to inhibit airway inflammation and promote airway relaxation may not fully address the complex biological and environmental factors that initiate AHR and acute exacerbation. In fact, studies have shown that O_3_ impairs the efficacy of glucocorticoid treatment in experimental models (8–10). Therefore, it is essential to understand the underlying mechanisms by which O_3_ triggers asthmatic AHR. To address this issue, our group recently developed a model of acute O_3_ exacerbation in mice with allergic airway inflammation. Using this model, we demonstrated that mice exposed to O_3_ in the setting of allergic inflammation had both increased AHR and changes in the innate and adaptive immune cell populations that comprise the inflammatory microenvironment within the lungs (11). In the present study, we build upon our prior work by utilizing single cell RNA sequencing to perform an unbiased analysis of immune cell transcriptomic changes in this niche to identify the cell populations responsible for O_3_ induced AHR during asthma exacerbations.

## Methods

### Mice

Male and female C57BL/6 J (stock number: 000664) mice (8-9 weeks old) were purchased from The Jackson Laboratory (Bay Harbor, ME) and allowed to rest for 1 week prior to experiments. Animals were housed in specific pathogen-free conditions within The Ohio State University Animal Care Facility (Columbus, OH). All animal experiments were performed in compliance with the U.S. Department of Health and Human Services Guide for the Care and Use of Laboratory Animals and were reviewed and approved by the Institutional Animal Care and Use Committee at The Ohio State University (Animal Use Protocol #2017A00000074-R2).

### Mixed allergen model of allergic asthma

The DRA triple-allergen mixture was made in PBS, with a final concentration of the following extracts: 0.83 mg/ml of house dust mite (*Dermatophagoides farina*), 8.33 mg/ml of short ragweed (*Ambrosia artenmisiifolia*), and 0.83 mg/ml of *Aspergillus fumigatus*, as previously described (11). All allergens were sourced from Stallergenes Greer. Mice were lightly anesthetized with 2-4% isoflurane before intranasal administration of 20 μl DRA mixture on experimental days 0, 5, 12, 14, and 17 (Fig. 1A). Under the same protocol, control mice were administered 20 μl PBS.

**Figure 1.**
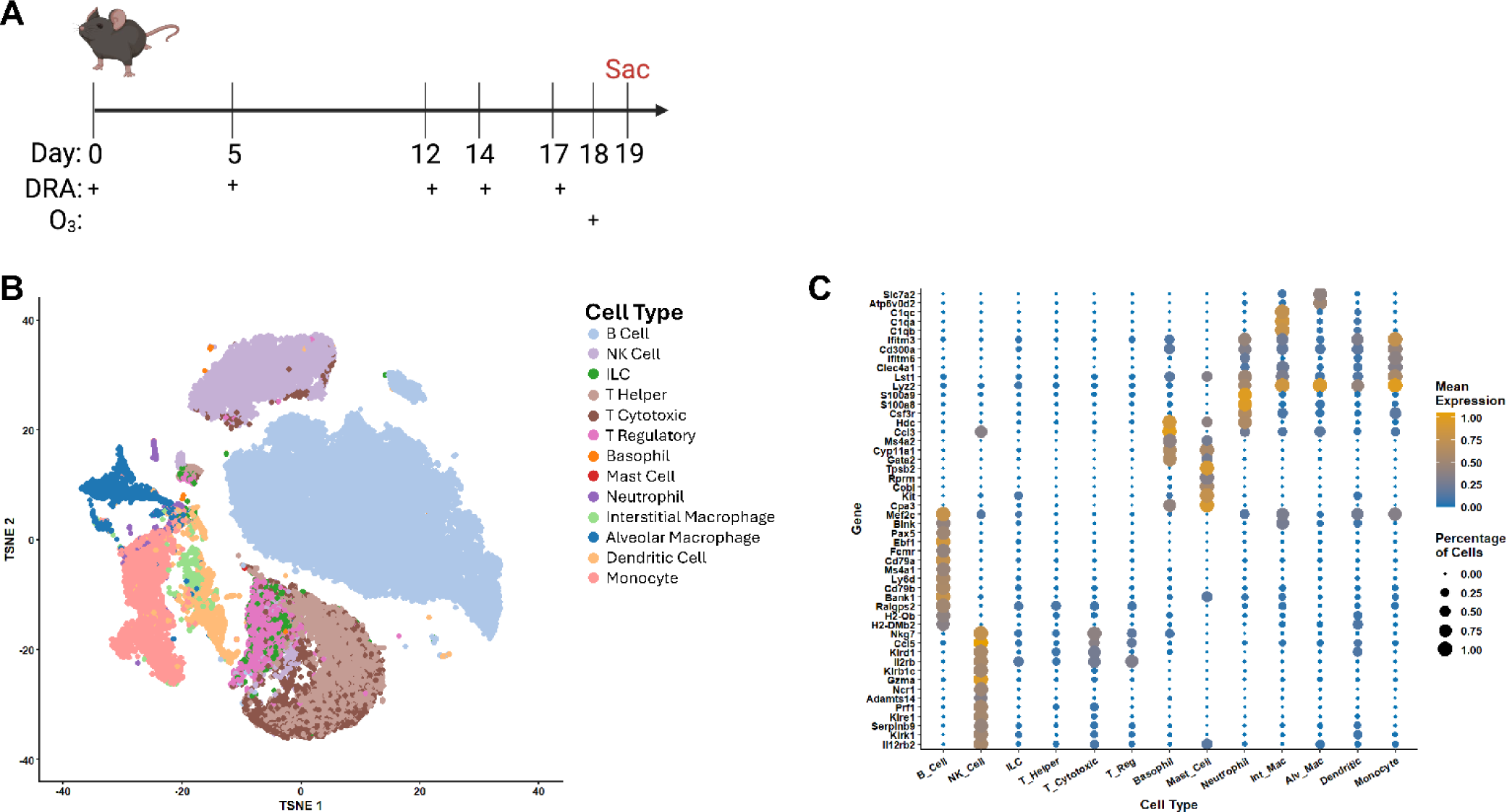
Identification of immune cell populations in the lungs of DRA+O_3_ mice using single cell RNA sequencing. As shown in (A), male and female C57BL/6 mice were intranasally sensitized and challenged with triple allergen mixture (DRA) five times over 17 days; 24 hr after the last DRA administration (day 18), mice were exposed to 2 ppm O_3_ for 3 hr; 24 hr after O_3_ (day 19) mice were sacrificed and tissues collected [image created with BioRender.com]. Whole lung tissue was digested using collagenase followed by live cell enrichment using FACS. Live cells were then processed for scRNA sequencing. (B) t-SNE overview of immune cell clusters identified in all 4 treatment groups (control, DRA, O_3_, DRA+O_3_). Clusters were annotated using previously validated marker genes. (C) Dot plot showing the average expression level (color) and percentage of cells expressing each gene (size) for the top 50 marker genes used in cluster identification. Dot plot showing expression of the top 50 marker genes based on level (color) and percentage of cells (size) for each cluster identified.

### Ozone exposure

On experimental day 18 (24 hr after the last DRA challenge), mice were exposed to room air (control) or O_3_ at an average of 2 ppm for 3 hr. This exposure paradigm was chosen to ensure O_3_-induced toxic effects in mice and is comparable to an O_3_ concentration of 400 ppb for humans (12). Whole-body exposures were performed using a custom-designed plexiglass chamber (13). O_3_ was generated by directing 100% oxygen over an ultraviolet light, which was then mixed with room air using an air pump and directed into the exposure chamber. O_3_ concentrations within the exposure chamber were monitored continuously (Photometric Ozone Analyzer – Model 400E, Teledyne Technologies) and adjusted accordingly. The temperature (average 21.6°C) and humidity (average dew point 17°C and relative humidity (80%) of the exposure chamber were also monitored.

### Euthanasia

Mice were euthanized on experimental day 19 (24 hr after O_3_ exposure / 48 hr after final DRA challenge) using pentobarbital (100 mg/kg I.P) or ketamine+xylazine (300 and 30 mg/kg, respectively) overdose.

### Bronchoalveolar lavage

0.9 ml of PBS was instilled into the bilateral lungs and withdrawn. This was repeated two times with the same fluid for cytokine analysis. The bronchoalveolar lavage (BAL) was centrifuged at ∼900*g* for 5 minutes to pellet any cells and the supernatant was collected for cytokine measurements. Four additional, separate washes (1 ml of PBS each) were performed, and the collected cells were pooled with those from the first aliquot for airway immune cell analysis. Total cell counts were determined by TC20™ Automated Cell Counter (BioRad).

### Single cell RNA sequencing sample preparation and library generation

Each treatment group was comprised of 4 male and 4 female mice. At the time of harvest, cells were isolated from whole lung tissue by collagenase digestion, as previously described (14). Lung cells from all males and all females were then pooled for each respective exposure group (*n* = 2 per exposure group; 1 pooled male sample and 1 pooled female sample). Live cells were then enriched using fluorescence activated cell sorting for DAPI^-^ cells. The live cell suspensions were then prepared for sequencing library generation by the Genomics Shared Resource at The Ohio State University Comprehensive Cancer Center according to the manufacturer’s protocol CG000315 Rev C for the Chromium Next GEM Single Cell 3’ Kit v3.1 (10X Genomics, Pleasanton, CA, USA).

### Single Cell Data processing

After demultiplexing, the filtered matrices where loaded into a Seurat (15) individually (min.cells = 0, min.genes=0, min.features=0), and percentage of mitochondrial content marked. To reduce noise in the data, cells with greater than 1% mitochondria and nFeatures greater than one where selected. Doublets (doublets_table.csv) were found and removed with scDblFinder (16). SCTransform (method = glmGamPoi, vars.to.regress = “perMito”) and Harmony (17) (group.by.vars = “orig.ident”, reduction = “pca”) was then performed to normalize and combine samples. The Azimuth (18) feature was used with LungMap (19) to annotate cell types. The mbkmeans (20) package was used to determine appropriate principle component analysis, dimensional reduction (PCA, UMAP, tSNE) was performed in Seurat and rendered using ggplot2 (21), and cell type annotation was compared to clustering to assure proper alignment.

### Pseudo-bulk RNA-Seq analysis

The 10x Genomics de-multiplexing pipeline retains many unwanted gene types; to reduce this unwanted noise, the Bioconductor (22) Mus.musculus package was used to annotate gene types, and non-protein coding genes removed from the Seurat object. Sequencing counts were aggregated, and differential analysis performed with EdgeR (23) as described in the EdgeR handbook (estimateDisp, qlmQLFit, qlmQLFTest) on cell types with greater than 1,000 cells.

### Transcriptional pathway enrichment analysis

Ingenuity Pathway Analysis (IPA; QIAGEN Inc.) was used to analyze all differentially expressed genes (DEG; FDR < .05, log(fold change) ≥ |1.0|) for transcriptional pathway enrichment analysis. The canonical and machine learning pathways presented in the main text were chosen based on biological relevance, statistical significance (*p* < .05), activation score (z-score ≥ |2.0|), and DEG overlap ratio. For AMs, DCs, and monocytes, a full list of IPA canonical pathways is provided in the supplemental text.

### Flow Cytometry

500,000 cells collected by BAL were stained with 1 μl LIVE/DEAD™ Fixable Dead Cell Stain (ThermoFisher) in 1 ml PBS for 15 minutes at room temperature. Cells were then incubated with Fc receptor block (anti-mouse CD16/32; BD biosciences) and subsequently stained with primary antibodies in 100 μl Flow Staining Buffer (Invitrogen) for 15 minutes at room temperature. Anti-mouse primary antibodies were used at the following dilutions to define immune cell populations (Fig. S3): Ly6G (1:100; Biolegend, 127628), CD45 (1:100; Biolegend, 103155), CD11b (1:100; Biolegend, 101241), CD8α (1:100; Biolegend, 100706), CD19 (1:100; Biolegend, 152407), CD64 (1:50; Biolegend, 139319), CD24 (1:100; Biolegend, 101824), CD4 (1:100; Biolegend, 100422), Siglec F (1:100, BD Pharmigen, 562680), CD11c (1:100; Biolegend, 117320), and CD3 (1:100; Biolegend, 100222). Spectral flow cytometry was performed using the Cytek® Northern Lights 3000 3-laser System in the Flow Cytometry Core at The Ohio State University Davis Heart and Lung Research Institute. All data analysis was performed in FlowJo version 10 software.

### Cytokine measurements

Cytokines in the BAL and plasma (Fig. S7) were measured using a Meso Scale Discovery Multiplex Assay according to manufacturer instructions. For the V-PLEX Mouse Cytokine 19-Plex Kit (IFNγ, IL-1β, IL-2, IL-4, IL-5, IL-6, KC/GRO, IL-10, TNFα, MCP-1, IL-33, MIP-1α, IP-10, MIP-2) BAL and plasma were diluted 1:4 and the extended sample incubation protocol was used as recommended. A custom U-PLEX Assay (mouse) was used to measure Eotaxin, IL-13, IL-17E/25, MCP-5/Ccl12, MIP-1β, MIP-3α, and VEGF-A in BAL and plasma; samples were diluted 1:2 and incubated for 1 hr.

### Statistics

For laboratory-based experiments, data sets were tested for normality via a Shapiro-Wilk test and statistical analyses were performed by comparison of means using a one-way ANOVA followed by Tukey’s *post hoc* testing. Statistical significance was defined as a probability of type I error occurring at less than 5% (*p* < 0.05). Graphics and analyses were performed in PRISM v 9 (GraphPad, San Diego, CA).

## Results

### Single cell RNA sequencing of lung immune cells after allergen and ozone exposure(s)

The DRA triple-allergen model (11) was used to induce allergic airway inflammation in wild-type mice. Both control and DRA sensitized mice were then exposed to O_3_ and sacrificed 24 hr post-exposure (Fig. 1A). The lungs were collected from all four treatment groups (control, DRA, O_3_, and DRA+O_3_) and processed for scRNA sequencing. A total of 49, 806 cell transcriptomes were profiled and passed quality control filtering; subsequent cell cluster annotation was performed using published studies (19). t-SNE dimensionality reduction technique was used to visualize all cell clusters, including structural and immune cells (Fig. S1A). The number of cells sequenced in each exposure group and a full list of marker genes can be found in the supplemental data (Table S1 & Fig. S1B). For immune cells, we captured both innate and adaptive immune cell populations (Fig. 1B). The top 50 marker genes used to identify immune cell populations are shown in Fig. 1C. Cell populations that were sufficiently abundant for subsequent differential expression analyses (>1,000 cells) included monocytes, alveolar macrophages (AMs), dendritic cells (DCs), T cells, B cells, and NK cells (Table S1).

### Alveolar macrophages from DRA+O_3_ exposed animals exhibit a unique transcriptional profile compared to DRA or O_3_ alone

AMs are the primary resident immune cells within the airways and play a critical role in regulating and directing inflammatory and functional responses to inhaled xenobiotics, including O_3_ (24). The transcriptional response of AMs to O_3_ exposure in the setting of pre-existing allergic inflammation has yet to be defined. When examining all immune cells, scRNA-seq analysis revealed that AMs displayed the greatest number of DEGs (Table S2). Therefore, we chose to perform subsequent analyses on these cells. AMs were identified in all four treatment groups (Fig. 2A, B) but differential expression analysis showed that AMs from the DRA+O_3_ group exhibited the greatest number of DEGs compared to controls (Fig. 2C, Table S2); a 300% and 900% increase compared to individual DRA and O_3_ exposures, respectively. Transcriptional changes in AMs from the DRA+O_3_ exposure were predominantly driven by the response to DRA (Fig. 2D); however, over half of the upregulated DEGs were unique to the DRA+O_3_ exposure group (Fig. 2D). AMs from DRA+O_3_ exposed mice demonstrated an alternative activation phenotype (increased *Arg1, Ccl24, Rnase2a*), which appeared to be driven by the DRA exposure (Fig. 2E). However, transcriptional pathway analysis revealed that after DRA+O_3_ exposure AMs had enhanced activation of interferon and interleukin signaling, and a relative inhibition of phagocytic and hypoxic pathways compared to DRA alone (Fig. 2F). In contrast, exposure to O_3_ alone enhanced gene expression in pathways related to sterol and cholesterol biosynthesis and cell cycle progression in AMs (Fig 2F). AMs from the 2-hit group also activated sterol and cholesterol biosynthesis, but without activating cell cycle-related pathways (Fig. 2F). Pathway analysis of AMs from DRA+O_3_ mice also revealed the unique activation of various pathways such as degradation of the extracellular matrix, differential regulation of cytokine production by IL-17A/F, cytokine storm signaling, response of EIF2AK1 to heme deficiency, and trafficking and processing of endosomal TLR (Fig. 2F). Taken together, these data highlight the divergence in biological processes involved in the AM response(s) to DRA-induced allergic inflammation compared to O_3_ alone, and further demonstrate that AMs developed unique transcriptomic profiles following exposure to DRA+O_3_ that were distinct from the individual exposures.

**Figure 2.**
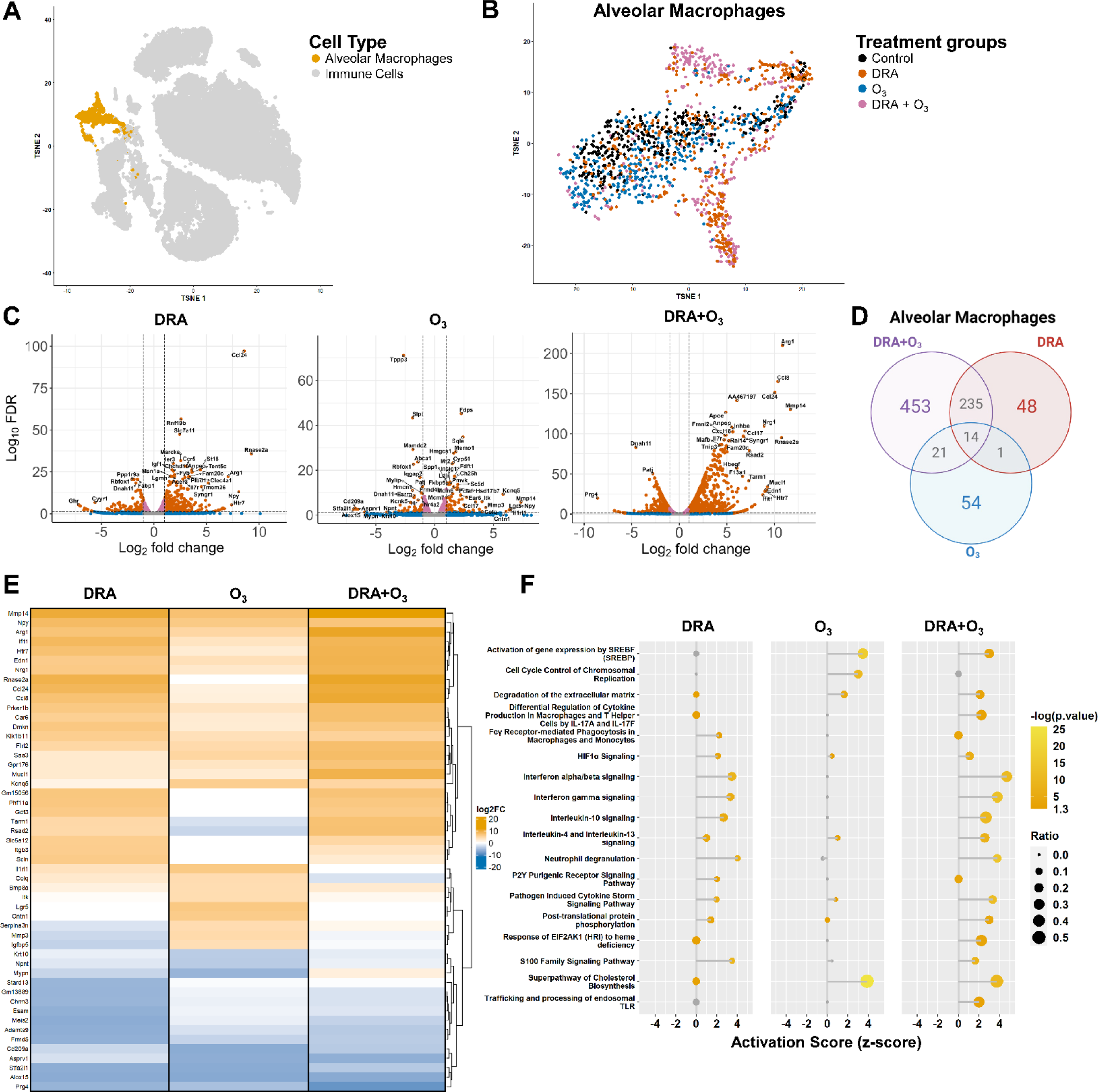
scRNA sequencing of AMs. (A) t-SNE plot of all immune cell clusters identified in the lungs of control, DRA, O_3_, and DRA+O_3_ mice, with AMs highlighted in orange. (B) t-SNE plot of AMs from all four groups; colors denote exposure(s) as indicated. (C) Volcano plots of differentially expressed genes (DEG) in AMs from DRA, O_3_, or DRA+O_3_ exposed mice (respectively) compared to AMs from the control group. Each dot represents one gene; “significant” defined as Log_10_(false discover rate [FDR]) < .05 and “relevant” mean expression level defined as Log_2_(fold change) > |1.0|, genes meeting both criteria considered differentially expressed (“relevant” orange dots above horizontal significance dashed line and outside vertical relevance dashed lines). (D) Venn diagram of the number of unique and overlapping DEG (upregulated only) in AMs from each treatment group compared to controls. (E) Heatmap displaying the top 50 DEGs in AMs from respective treatment groups compared to controls. (F) Dot plot displaying transcriptional pathway enrichment analysis in AMs from respective treatment groups compared to control AMs. The x-axis denotes respective pathway activation scores (z-score ≤ - 2 considered inhibited and ≥ 2 considered activated) and dot color corresponds to -log(p-value); -log(0.05) = 1.3, therefore pathways with -log(p-value) ≤ 1.3 are considered non-significant, as indicated by grey dot color. The dot size is proportionate to DEG overlap ratio [DEG in data set/total genes in pathway].

### O_3_ exposure in the setting of allergic inflammation leads to the presence of a unique monocyte population within the lungs

Circulating monocytes survey the lung during homeostasis (25) but can also be recruited during inflammation and injury to serve as precursors for resident macrophage populations such as AMs. Initial transcriptomic analysis revealed that there was a unique population of lung monocytes in the 2-hit group (Fig. 3B). Compared to controls, monocytes from the lungs of DRA exposed mice only had 35 DEGs, while O_3_ and DRA+O_3_ exposure induced differential expression of 179 and 91 genes, respectively (Fig. 3C, Table S2). The transcriptional changes in monocytes from the O_3_ exposure group were predominantly downregulation of gene expression (Fig. 3C, Table S2), so >90% of the DEGs upregulated in DRA+O_3_ monocytes were unique to the 2-hit exposure (Fig. 3D). The DEGs with the greatest increase in expression in DRA monocytes relative to controls were *Clca1* (6.485 fold increase) which promotes mucus production (26) and *Reg3g* (6.553 fold increase) which has been reported to suppress allergic airway inflammation (27) (Fig. 3E). In DRA mice subsequently exposed to O_3_, these genes were no longer increased; instead, DRA+O_3_ monocytes showed high expression of chemokine *Cxcl3* (Fig. 3E). Pathway analysis of significant DEGs (Fig. 3C) showed that there were few differentially regulated pathways in lung monocytes following DRA and O_3_ exposures compared to controls (Fig. 3F). After O_3_ exposure alone, monocytes in the lungs showed an inhibition of interferon signaling (Fig. 3F). Alternatively, exposure to both DRA+O_3_ caused activation of pro-inflammatory pathways in lung monocytes, such as cytokine storm and S100 family signaling, as well as GPCR signaling and phagosome formation (Fig. 3F). These data demonstrate that transcriptional pathways in lung monocytes differ in the DRA+ O_3_ model compared to uninjured controls or either exposure alone.

**Figure 3.**
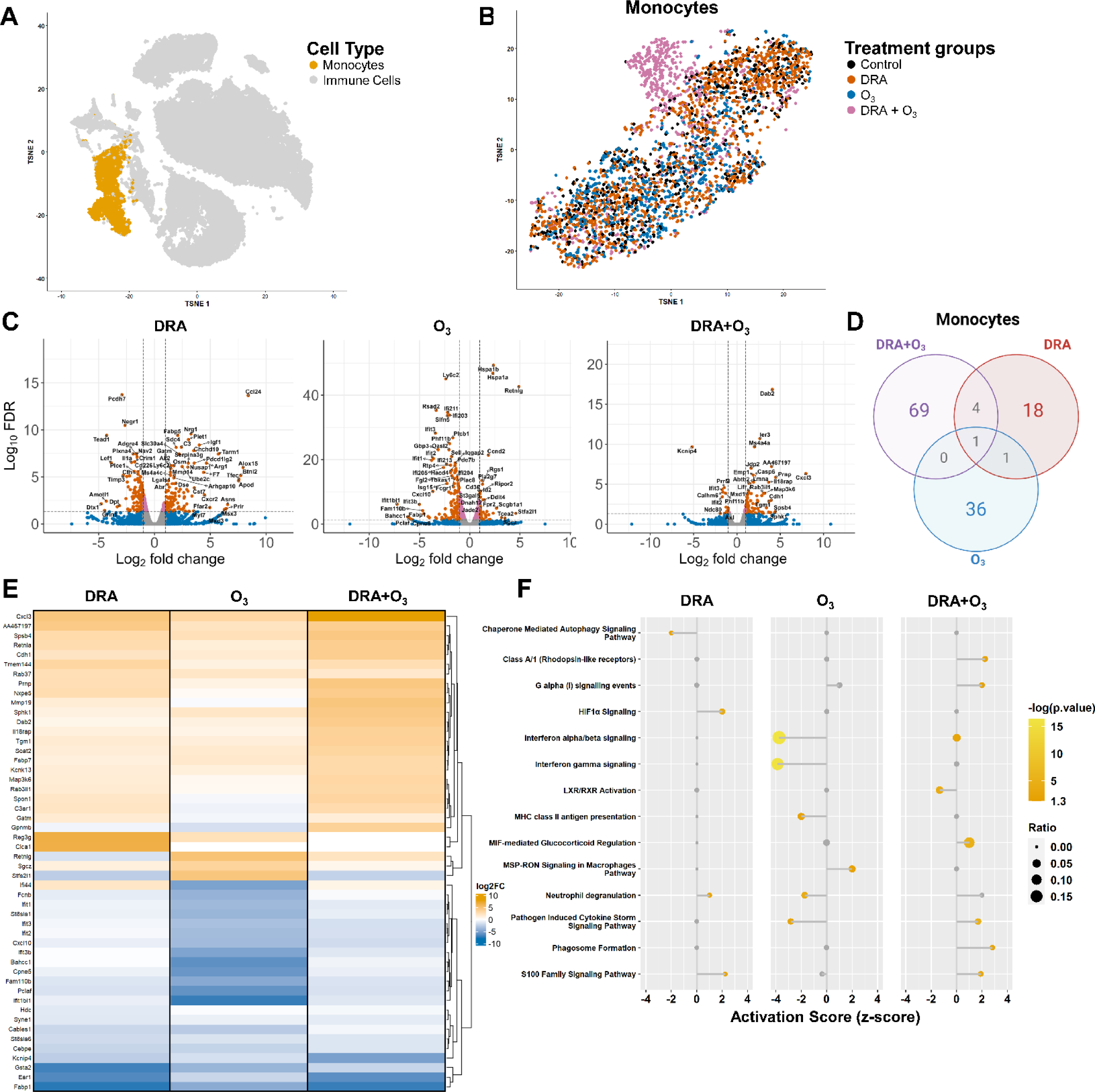
scRNA sequencing of monocytes present in the lungs of DRA, O_3_, and DRA+O_3_ exposed mice. (A) t-SNE plot of all immune cell clusters identified in the lungs of control, DRA, O_3_, and DRA+O_3_ mice, with monocytes highlighted in orange. (B) t-SNE plot of monocytes from all four groups; colors denote exposure(s) as indicated. (C) Volcano plots of differentially expressed genes (DEG) in monocytes from DRA, O_3_, or DRA+O_3_ exposed mice (respectively) compared to monocytes from the control group. Each dot represents one gene; “significant” defined as Log_10_(false discover rate [FDR]) < .05 and “relevant” mean expression level defined as Log_2_(fold change) > |1.0|, genes meeting both criteria considered differentially expressed (“relevant” red dots above horizontal significance dashed line and outside vertical relevance dashed lines). (D) Venn diagram of the number of unique and overlapping DEG (upregulated only) in monocytes from each treatment group compared to controls. (E) Heatmap displaying the top 50 DEGs in monocytes from respective treatment groups compared to controls. (F) Dot plot displaying transcriptional pathway enrichment analysis in monocytes from respective treatment groups compared to control monocytes. The x-axis denotes respective pathway activation scores (z-score ≤ -2 considered inhibited and ≥ 2 considered activated) and dot color corresponds to -log(p-value); - log(0.05) = 1.3, therefore pathways with -log(p-value) ≤ 1.3 are considered non-significant, as indicated by grey dot color. The dot size is proportionate to DEG overlap ratio [DEG in data set/total genes in pathway].

### DRA-mediated changes in dendritic cell phenotype are maintained after O_3_ exposure

DCs are professional phagocytes and antigen presenting cells that have been shown to play an important role in allergic lung inflammation. Differential expression analysis demonstrated that these cells had the second highest number of DEGs in response to DRA+O_3_ (Table S2); however, in contrast to AMs and monocytes, tSNE plots of DCs did not indicate any potentially unique transcriptional subsets between treatment groups (Fig. S2B). Of the 3 exposure groups, DCs from the 2-hit group had the greatest number of DEG compared to controls (Fig. S2C, Table S2). Approximately half of the upregulated DEG in DCs from the DRA+O_3_ group were unique to the 2-hit model and ∼25% overlapped with DRA alone (Fig. S2D). Transcriptional pathway analysis revealed minimal changes after O_3_ exposure (Fig. S2F). The majority of pathways activated following DRA, with or without subsequent O_3_ exposure, were related to cellular division, antigen presentation, and leukocyte extravasation (Fig. S2F). Interestingly, O_3_ as a second hit appeared to impair the DRA-mediated activation of cell cycle pathways in DCs, but increased MHCII antigen presentation (Fig. S2F). These results suggest that DRA exposure induces activation of cell cycle pathways in DCs in the lungs and subsequent O_3_ exposure impairs this response.

### Allergic inflammation causes recruitment of macrophage subtypes into the airways but impairs neutrophil recruitment following ozone

Transcriptional analysis revealed differences in gene expression of immune cells following DRA and/or O_3_ exposure. Previously, we performed flow cytometry assessment of immune cell populations in whole lung tissue (11). However, since the current study suggested a predominant role of AMs in the response to the 2-hit DRA+O_3_ model, we sought to comprehensively characterize the airspace niche, where the AMs reside, by analyzing the immune cell populations present in bronchoalveolar lavage (BAL). Compared to controls, total BAL cell counts revealed significant immune cell infiltration in DRA-treated mice with or without O_3_ exposure, but not in response to O_3_ alone (Fig. 4A). Flow cytometry was used to analyze the immune cell populations from the BAL (Fig. S3, S4). In control mice, tissue-resident (TR)-AMs made up 90% of the immune cells present in the airspace (Fig. 4B). This proportion decreased following all exposures, as other immune cells were recruited to the lung (Fig. 4B). At the time point these samples were collected, there was no difference in the absolute number of TR-AMs in any of the experimental groups (Fig. 4C). Exposure to DRA with or without O_3_ induced recruitment of monocyte-derived (Mo)-AMs, interstitial macrophages (IMs), and DCs into the airways, but this was not observed in mice exposed to O_3_ alone (Fig. 4D-F). Granulocytes were absent in the airspace of control mice, but DRA exposure induced significant eosinophilia that was not affected by subsequent O_3_ exposure (Fig. 4G). O_3_ exposed mice showed a significant increase in neutrophil numbers compared to controls; however, O_3_-induced neutrophil recruitment was impaired in the setting of allergic inflammation in the DRA+O_3_ group (Fig. 4H). Adaptive immune cells were mostly absent in the BAL of control mice, but DRA exposure induced a significant increase in the number of B cells, CD8^+^ T cells, and CD4^+^ T cells that was unaffected by subsequent O_3_ exposure (Fig. 4I-K). These data highlight the extensive differences in the immune cells present in the airspace niche of normal and DRA-exposed mice that can alter the microenvironment and consequently the AM response to inhaled O_3_.

**Figure 4.**
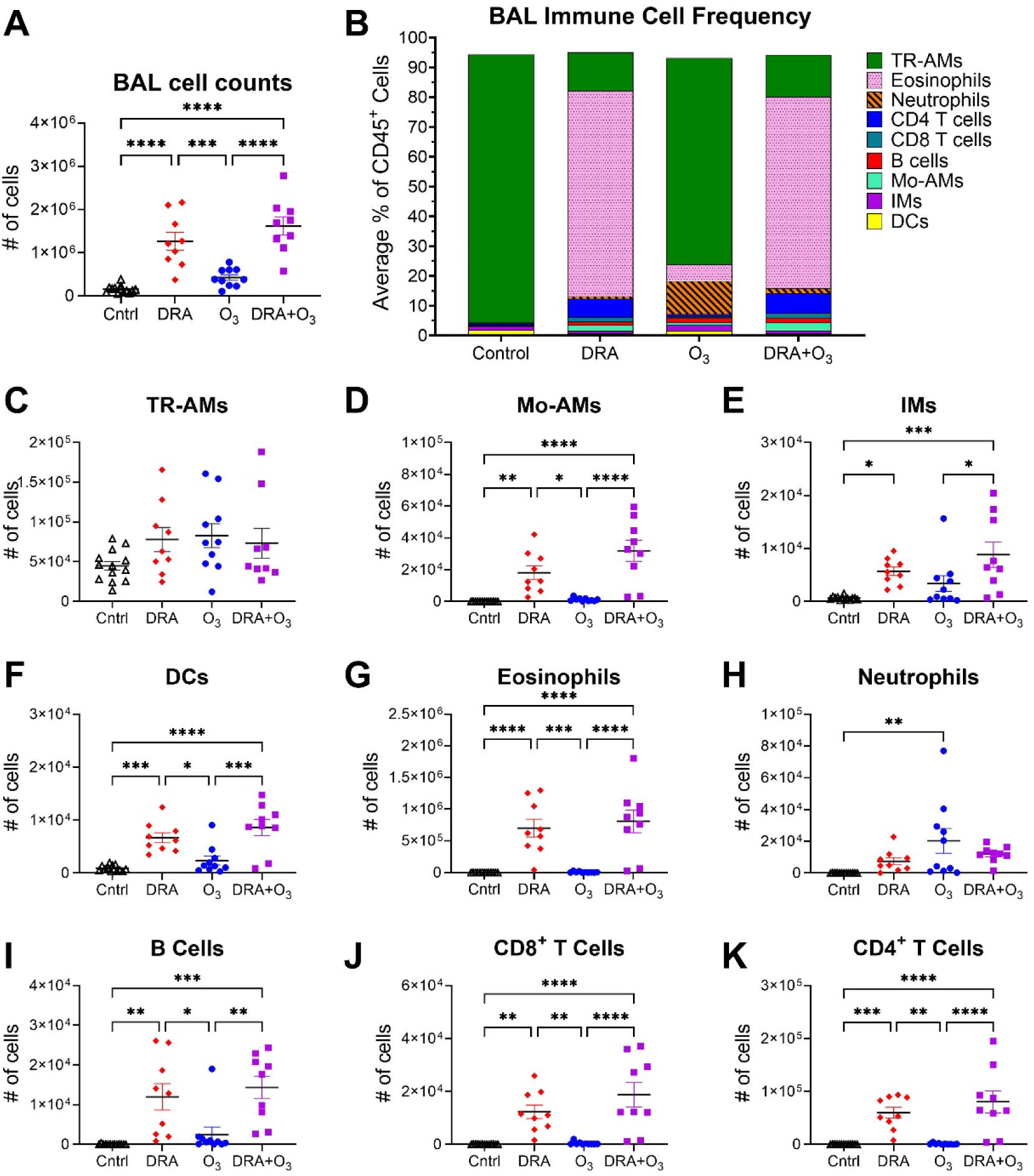
Airspace inflammatory cells in DRA, O_3_, and DRA+O_3_ exposed mice. DRA triple allergen administration and O_3_ exposure was performed in male and female C57BL/6 mice as previously described (Figure 1A). Bronchoalveolar lavage (BAL) was performed and cells within the airspaces collected for (A) total cell counts, and flow cytometry used to determine (B) frequency and (C-K) total number of immune cell populations in the BAL. *n* = 9-13; \**p* < .05, \*\**p* < .01, \*\*\**p* < .001 as indicated; statistical significance analyzed using one-way ANOVA followed by Tukey’s *post hoc* testing. Data in panels A and C presented as *mean ± SEM*, data in panel B presented as *mean* only. AM, alveolar macrophage; TR, tissue-resident; Mo, monocyte-derived; AM, alveolar macrophage; IM, interstitial macrophage; DC, dendritic cell.

### Pre-existing allergic inflammation alters the inflammatory mediators present within the airspaces following O_3_ exposure

To further characterize the inflammatory signaling within the airspace, we measured cytokine levels in the BAL fluid (Fig. 5, S5, S6). Type 2 cytokines IL-4 and IL-13 were increased over controls in both DRA and DRA+O_3_ groups (Fig. 5A, B). Although DRA-mediated eosinophilia in the BAL was unaffected by O_3_ (Fig. 4G), eosinophil maturation and chemotactic factors, IL-5 and CCL11 (Eotaxin), were increased in DRA+O_3_ compared to DRA alone (Fig. 5C, D). Despite reduced neutrophil numbers in the BAL of DRA+O_3_ mice compared to O_3_ alone (Fig. 4H), neutrophil chemoattractants CXCL1 (KC/GRO) and CXCL2 (MIP-2) were increased over controls after DRA+O_3_ exposure; in fact, CXCL2 was significantly higher in the DRA+O_3_ group compared to O_3_ alone (Fig. 5 E, F). CCL2, also known as monocyte chemoattractant protein (MCP)-1, was only significantly increased after DRA+O_3_ (Fig. 5 H). Additional chemokines and pro-inflammatory cytokines that were increased in the DRA-exposed groups (DRA and DRA+O_3_), but not after O_3_ only exposure, included: CXCL10 (IP-10), CCL3 (MIP-1α), CCL4 (MIP-1β), IL-1β, and TNFα (Fig. 5G, I, J, L, N). Interestingly, chemokine CCL20 (MIP-3α) was significantly increased over controls in the individual exposure groups, but there was no difference following the 2-hit exposure (Fig. 5K). The increase in the acute inflammatory cytokine IL-6 was driven by O_3_ exposure (Fig. 5M). Finally, expression of VEGF-A was only increased over controls following O_3_ exposure; induction of this factor by O_3_ was impaired by allergic inflammation in the DRA+ O_3_ group (Fig. 5O). We hypothesize that these changes in cytokine and chemokines within the airspace were due to the cells present in this microenvironment, because there were only limited changes observed in serum of these mice (Fig. S7). These data demonstrate that the inflammatory milieu within the airspace after DRA+O_3_ is primarily driven by the response to DRA allergens, but pre-existing DRA-induced inflammation alters the microenvironment during subsequent O_3_ exposure (DRA+O_3_).

**Figure 5.**
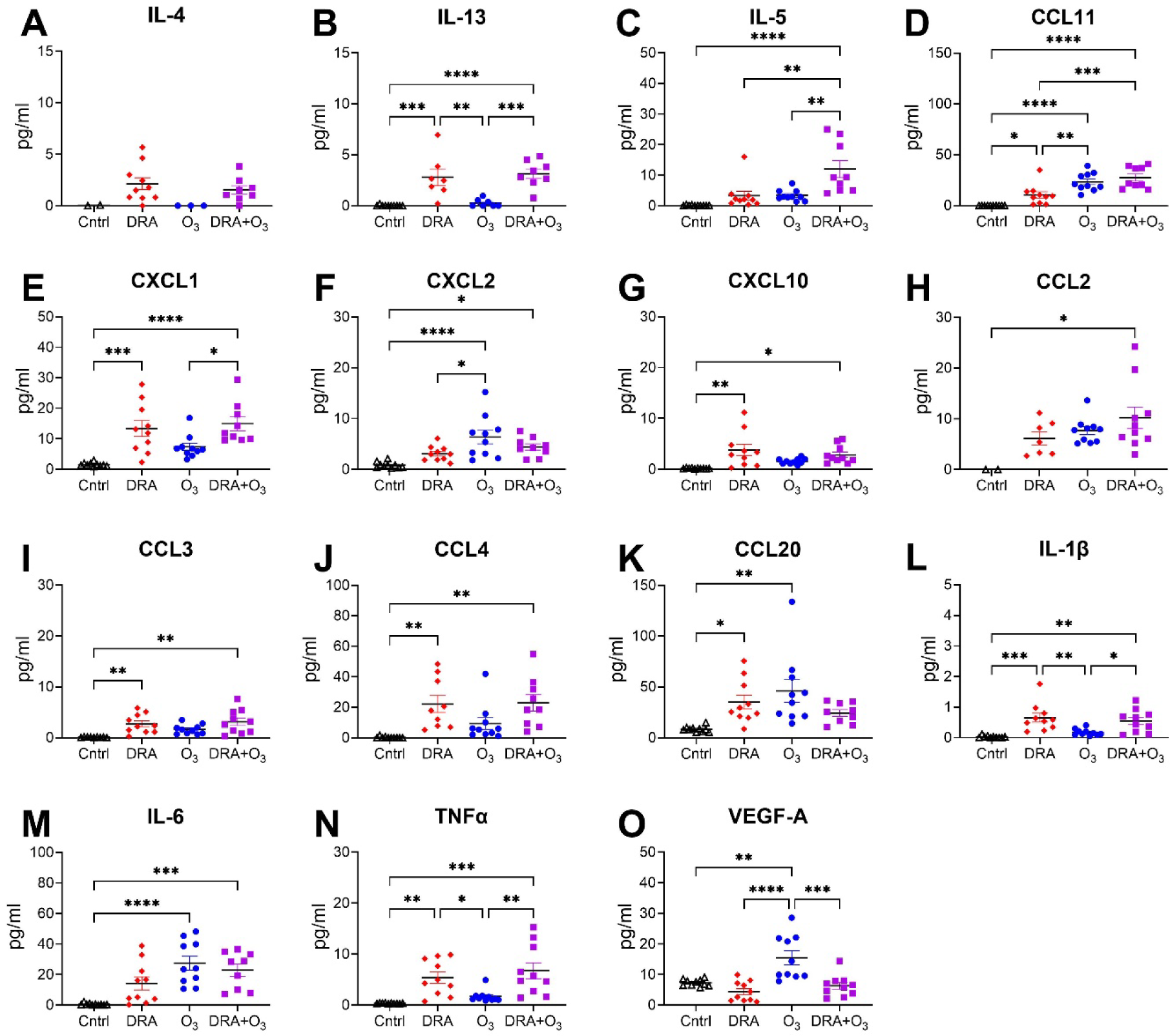
Airspace cytokine levels in DRA, O_3_, and DRA+O_3_ exposed mice. As shown in Figure 1A, male and female mice were intranasally sensitized and challenged with triple allergen mixture (DRA) five times over 17 days; 24 hr after the last DRA administration, mice were exposed to 2 ppm O_3_ for 3 hr; 24 hr after O_3_ concentrated bronchoalveolar lavage fluid (BALF) was collected for airspace measurement of cytokines and chemokines. All data presented as *mean ± SEM*; *n* = 9-10; \**p* < .05, \*\**p* < .01, \*\*\**p* < .001 as indicated; statistical significance analyzed using one-way ANOVA followed by Tukey’s *post hoc* testing.

**Figure 6.**
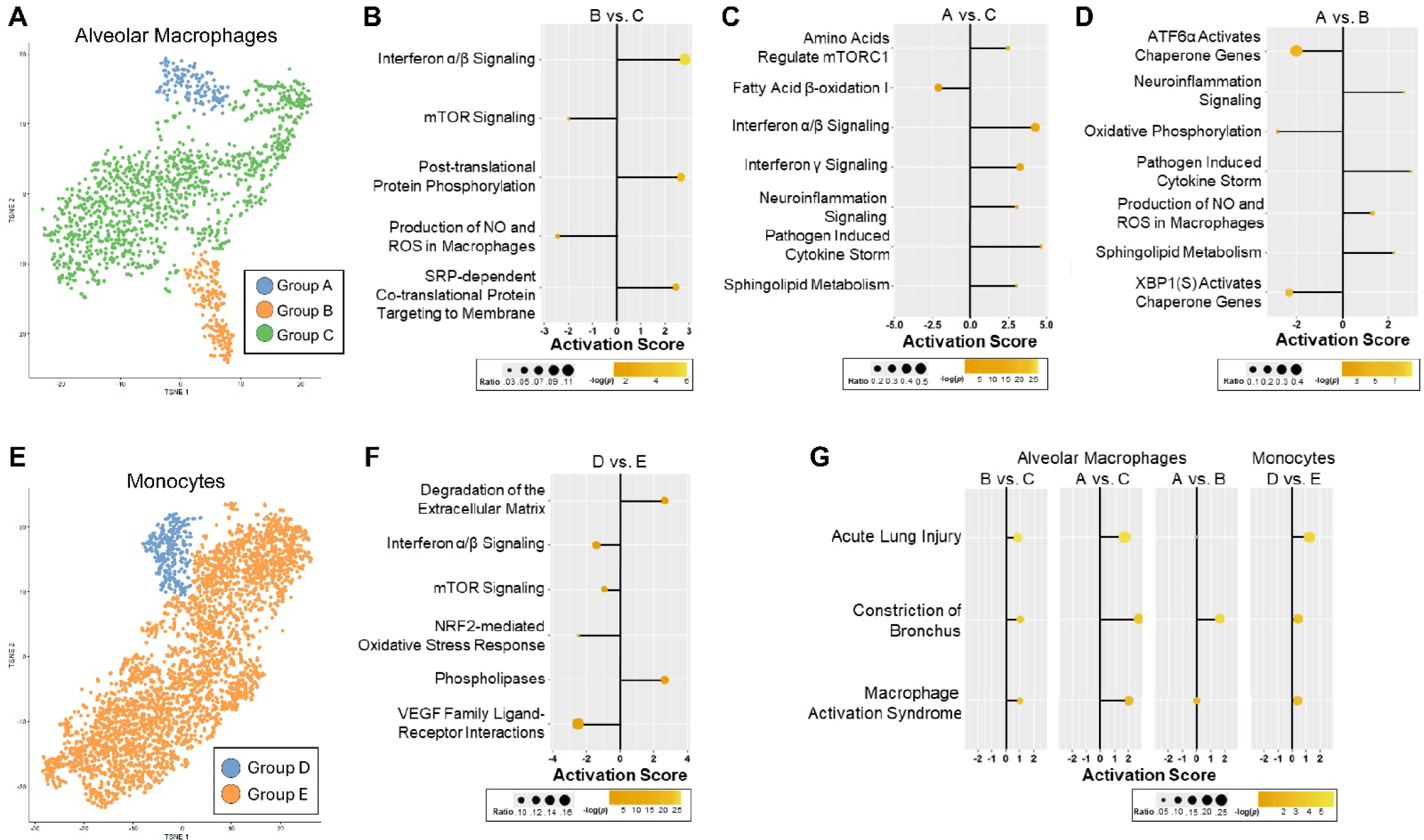
Evaluation of AM and monocyte subpopulations unique to DRA+O_3_ exposure. (A) t-SNE plot displaying AMs from all 4 exposures (Figure 2B); colors denote unique transcriptional groups manually defined for differential expression analysis. Group “A” is predominantly AMs from DRA+O_3_, “B” is comprised of AMs from DRA and DRA+O_3_ groups, and “C” contains AMs from all 4 exposures but dominated by control and O_3_ alone. (B-D) Canonical pathway enrichment analysis (IPA software) of DEGs between AM groups, as indicated. (C) t-SNE plot displaying monocytes from all 4 exposure groups (Figure 3B); colors denote unique transcriptional groups manually defined for differential expression analysis. Group “A” contains monocytes exclusively from the DRA+O_3_ exposure paradigm, and group “B” is a mixture of cells from all 4 exposures. (F) Canonical pathway enrichment analysis (IPA software) of DEGs between monocyte groups “D” and “E”. (G) Machine Learning Disease Pathways based on DEGs between AM or monocyte groups, as indicated. For all dot plots the x-axis denotes respective pathway activation scores (z-score ≤ -2 considered inhibited and ≥ 2 considered activated) and dot color corresponds to -log(p-value); -log(0.05) = 1.3, therefore pathways with -log(p-value) ≤ 1.3 are considered non-significant, as indicated by grey dot color. The dot size is proportionate to DEG overlap ratio [DEG in data set/total genes in pathway].

### A transcriptionally distinct AM subpopulation unique to the DRA+O_3_ exposure activates disease pathways related to AHR

We further examined AM and monocyte transcriptional subpopulations by performing a manual sub-setting based on dimensionality reduction t-SNE plots. Using this technique, there were 3 potentially distinct AM subsets (Fig. 6A). AM subset “A” was comprised of cells primarily from the DRA+O_3_ exposure group, subset “B” contained AMs from both DRA and DRA+O_3_ exposures, and “C” was made up of AMs from all 4 exposure groups but predominantly from control and O_3_ exposures (Fig. 2B, 6A). Pathway analysis of subset “B” revealed enrichment of genes involved in interferon α/β signaling, protein targeting to the membrane, and posttranslational phosphorylation pathways, but downregulation of genes related to mTOR signaling and nitric oxide (NO) and reactive oxygen species (ROS) production (Fig. 6B). In AM subset “A” (DRA+O_3_) mTORC1 signaling was activated compared to AMs in “C” (control + O_3_) (Fig. 6C), which is consistent with the activation of sphingolipid metabolism pathway and inhibition of fatty acid β-oxidation pathways also observed in “A” (Fig. 6C). AM subset “A” was also enriched for genes involved in multiple pathways related to inflammatory signaling (Fig. 6C). Finally, comparing AM subset “A” (DRA+O_3_) to “B” (DRA±O_3_) allowed us to examine the unique transcriptional changes driven by O_3_ as a second hit following allergic inflammation (Fig. 6D). Transcriptional pathway analysis revealed a relative inhibition of oxidative phosphorylation and chaperone gene pathways related to ER stress, as well as upregulation of genes involved in NO and ROS synthesis pathways, sphingolipid metabolism, and inflammatory pathways related to neuroinflammation and cytokine storm signaling (Fig. 6D). In contrast to the subgroup analysis of AMs, there were only 2 subgroups identified on dimensionality reduced monocyte t-SNE plots (Fig. 6E). The cells in subset “D” were only from the DRA+O_3_ exposure group, while “E” was comprised of monocytes from all 4 exposures (Fig. 3B, 6E). Compared to all other monocytes, the DRA+O_3_-specific subset (“D”) had decreased gene expression related to mTOR signaling which was reflected in the activation of pathways related to degradative phospholipases and inhibition of VEGF family interactions and the NRF2 oxidative stress response (28, 29)(Fig. 6F).

To further characterize the contributions of these subsets to the functional AHR response to O_3_, we utilized IPA’s Machine Learning evaluation of association with disease processes. These analyses indicated that following DRA treatment, AM subsets “A” and “B” activated processes related to macrophage activation syndrome and acute lung injury, as compared to subset “C” (Fig. 6G). Interestingly, subset “A” (overrepresentation by DRA+O_3_) was most significantly involved in processes associated with constriction of bronchus (Fig. 6G), corresponding with the enhanced functional AHR response (increased airway resistance) previously observed in DRA+O_3_ mice (11). Comparison of monocyte subsets did not show significant activation (z-score < 2.0) of any of the presented disease pathways (Fig. 6G, panel 4). Taken together, these results point towards a unique transcriptional subset of AMs that contributes to not only inflammatory signaling (Fig. 6C, D) but functional AHR outcomes following DRA+O_3_ exposure (Fig. 6G).

## Discussion

More than 350 million people worldwide live with asthma every day (1), but exposure to various triggers, such as allergens and air pollution can cause exacerbation or worsening of asthma symptoms and lung function. Previously, we established a murine model of DRA-induced allergic lung inflammation followed by acute O_3_ exposure to mimic the human asthmatic AHR response for mechanistic investigation (11). In the present study, we build upon this work by performing scRNA sequencing on lung cells of mice exposed to DRA, O_3_, or DRA+O_3_ to gather unbiased insights into the cell types involved in the functional response to inhaled O_3_ in the context of allergic lung inflammation. We were able to detect changes in both the innate and adaptive immune cells within the lung, but the most significant findings were the transcriptional changes observed in AMs in response to DRA+O_3_ (Fig. 2, Table S1). Upon further analysis, there were transcriptionally distinct groups of both AMs and monocytes that were unique to the DRA+O_3_ group (Fig. 6). Pathways that were upregulated in these subsets of cells include production of cytokines and oxidants, lipid metabolism, and bronchial constriction. Taken together these data highlight an important role for macrophage activation by O_3_ in the allergic lung niche.

O_3_ inhalation causes lung injury and since AMs are the primary immune cell present within the airways, they play an essential role in protecting the host against inflammatory insults. In the present study, AMs demonstrated minimal transcriptional changes in response to O_3_ alone, except for activation of cholesterol biosynthesis and cell cycle pathways (Fig. 2F). AMs from the DRA+O_3_ exposure group also upregulated gene pathways related to cholesterol biosynthesis, but in the absence of cell cycle pathway activation (Fig. 2F). Targeted analysis of unique AM subpopulations revealed that a subgroup of AM comprised cells mostly from DRA+O_3_ (group “A”, Fig 6) had transcriptional changes in canonical pathways related to lipid metabolism, oxidative phosphorylation, and inflammatory signaling (Fig. 6C, D). Interestingly, altered glycosphingolipid metabolism has been previously associated with the allergic AHR response to O_3_ (30) and individual sphingolipids have been shown to enhance airway smooth muscle (ASM) cell proliferation and AHR (31). Furthermore, AMs and monocytes from DRA+O_3_ exposed mice, but not DRA alone, activated transcriptional pathways related to degradation of the extracellular matrix (Fig. 2F, 6F) including the upregulation of matrix metalloprotease (MMP) gene expression (Fig. 2E, 3E). In accordance with these results, AMs from people with asthma have been shown to release higher levels of MMPs, which is associated with increased airway thickness and a faster decline in FEV_1_ (32). These changes in AM phenotype following the 2-hit exposure demonstrate that the immune response to inhaled O_3_ is different in asthmatic lungs compared to normal lungs.

Airway macrophages are comprised of cells from two different ontogenies: fetal yolk sac derived self-renewing TR-AMs and recruited Mo-AMs differentiated from circulating monocytes originating from the bone marrow (33). Following exposure to DRA there was recruitment of Mo-AMs into the airspaces, which remained present during subsequent exposure to O_3_; but these recruited Mo-AMs were absent in mice challenged with O_3_ alone (Fig. 4D). TR-AMs are considered protective against allergic inflammation and essential for resolution of O_3_-induced inflammation (34, 35) and a recent study also showed a population of recruited CX_3_CR ^+^ AMs facilitate the resolution of eosinophilic inflammation in the DRA model (36). However, in contrast to these protective roles, recruited monocytes/Mo-AMs are reported to drive lung inflammation and injury in a variety of models (33, 37–41), including allergic inflammation and allergen-induced AHR (34, 42, 43). In accordance with the present findings (Fig. 4D), a lineage tracing study recently demonstrated that there is no recruitment of Mo-AMs in healthy mice or humans exposed to O_3_ alone (35). Consequently, the influence of Mo-AMs in the asthmatic AHR response to O_3_ has yet to be directly investigated.

There are other immune cells present within the allergic asthmatic lung that can also regulate downstream signaling responses to O_3_ exposure. Neutrophils are recruited to the airways following O_3_ inhalation (Fig. 4) and airway neutrophilia is linked to corticosteroid resistance and severe asthma (44). Reports on the role of neutrophils in AHR have been mixed; some studies report a direct role in AHR (45–47) while others do not (48, 49). Neutrophils are recruited via CXCL1 and IL-6, both of which were elevated in the BAL following DRA+O_3_ (Fig. 5E, M). There was no significant difference in airspace neutrophilia between O_3_ and DRA+O_3_ exposure groups (Fig. 4H), however, significant differences were observed when the data was analyzed by sex (Fig. S4C). Macrophages present within the lung interstitium can also modulate lung inflammation and remodeling. Previous reports demonstrate that IL-10 production by IMs prevents neutrophilic asthma (50), but that LPS-mediated replacement of embryonically-derived IMs with bone marrow-derived IMs reduces BAL IL-10 and AHR in response to subsequent allergen challenge (51). The current study reports increased IM numbers in the BAL of DRA-exposed mice (Fig. 4E). But, we have previously demonstrated that O_3_ attenuates DRA-induced increases in IM numbers and IL-10 levels in the lung tissue of male mice (11), which corresponds with the increased neutrophils numbers observed in the BAL of DRA+O_3_ exposed males (Fig. S4C). It is therefore possible that O_3_ changes IM function in a deleterious way, however, further work is needed to understand the role of these cells in regulating the AHR response.

Although inflammatory cells regulate various aspects of lung injury and allergic inflammation, bronchoconstriction and acute shortness of breath due AHR are mediated by the contraction of airway smooth muscle (ASM) cells. ASM contraction is induced by cellular mediators such as histamines, trypase, and eicosanoids, as well as acetylcholine released from neurons (5). Immune cell inflammatory signaling influences this contractile response and contributes to the hypercontractility observed in asthmatic AHR. Previous studies have identified TNFα, IFNγ, IL-13, and IL-1β as inflammatory cytokines that alter ASM phenotype through mechanisms such as enhanced proliferation and contractile force, corticosteroid resistance, and increased expression of adhesion molecules that facilitate ASM interactions with immune cells (52, 53). Most of these cytokines were significantly increased in the BAL of DRA+O_3_ mice compared to O_3_ alone (Fig. 5). Type 2 cytokines IL-4 and IL-5 were also elevated in the BAL of DRA and DRA+O_3_ exposed mice (Fig. 5 A, C); IL-4 directly promotes AHR in human bronchi and hypercontractility in ASM cells (54), while IL-5 contributes indirectly to altered ASM function through the recruitment of eosinophils, which can directly stimulate ASM contraction and increase acetylcholine signaling from lung neurons, resulting in AHR (55, 56). In the present study, IL-5 levels were higher in the 2-hit group compared to either exposure alone (Fig. 5C), but eosinophil numbers in the BAL were not different between DRA and DRA+O_3_ groups (Fig. 4G) and were reduced in the lung tissue of DRA+O_3_ compared to DRA alone (11). These data imply that differences in eosinophilia do not explain the increased AHR following DRA+O_3_ compared to DRA alone (11) at this time point.

Research examining ozone exacerbation of asthma in humans has been primarily epidemiological and/or descriptive in nature, not necessarily mechanistic. However, our previous study (11) recapitulates extensive human data demonstrating that short-term ozone exposure increases risk of asthma exacerbations (defined by lung function) for several days post-exposure (57). Herein we also show that our model aligns with human studies reporting O_3_ causes an increase in the number of airway immune cells, including macrophages and neutrophils (Fig. 4) (7, 44). Our model uses a single O_3_ exposure while asthmatic patients can experience sustained exposure(s) over multiple days, which may affect the various inflammatory cells and signals present in the lungs following O_3_ inhalation. However, the cells and inflammatory mediators necessary to exacerbate the functional AHR response to O_3_ were present and therefore this model and time point prove meaningful in elucidating the cellular mechanisms of asthmatic AHR. Another limitation of the present study was the number of cells required for pseudo-bulk sequencing analysis. As such, there were insufficient cell numbers to examine transcriptional changes in granulocytes, IMs, or structural cells such as ASM. Thus, future work will aim to investigate the interactions between various granulocyte and macrophage populations with ASM in the 2-hit model. Additionally, cell number requirements necessitated pooling of data from male and female mice and limited the ability to determine whether any sex differences in transcriptional changes following DRA and/or O_3_ were present. However, sex-based BAL flow cytometry and cytokine assessments can be found in the supplemental data (Fig. S4, S6).

In summary, asthma is responsible for over 1,300 deaths per day (1) and ∼40% of asthmatics in the US have had an acute exacerbation (“asthma attack”) in the past year (58). Furthermore, as climate change continues to worsen and ambient O_3_ levels continue to rise (2), vulnerable populations will become increasingly at risk of experiencing the deleterious consequences on lung function. Therefore, expanding asthma research to focus on mechanisms of non-allergen triggers of AHR, especially pervasive environmental pollutants such as O_3_, is essential to mitigating disease burden. The relationship between asthmatic AHR and granulocyte/inflammatory cell presence in the airways is still being debated (59, 60), but there is a large body of work that points towards a central role for AMs in regulating this process (53). Nevertheless, there is still a critical gap of knowledge regarding the direct interactions between AMs and ASM cells. The results of the present study emphasize the significant role AMs play in the response to inhaled O_3_ and emphasize the need for further research investigating the cross talk between macrophage subpopulations and the structural cells responsible for AHR-mediated bronchoconstriction.

## Author contributions

- Jess Ray: data acquisition, analysis, and interpretation, writing – original draft, writing – review & editing
- Josh Walum: data analysis, methodology, writing – review & editing
- Daria Jelic: data acquisition and interpretation, writing – review & editing
- Ryelie Barnes: data acquisition, writing – review & editing
- Ian Bentley: data acquisition, writing – review & editing
- Rodney D. Britt Jr.: conceptualization, methodology, data interpretation, writing – review & editing
- Joshua A. Englert: conceptualization, methodology, writing – original draft, writing – review & editing, supervision, project administration, funding acquisition.
- Megan N. Ballinger: conceptualization, methodology, writing – original draft, writing – review & editing, supervision, project administration, funding acquisition.

## Acknowledgements

The authors would like to thank Dr. Rama Mallampalli, MD (The Ohio State University, Department of Internal Medicine) for generously allowing us to use their laboratory’s Meso Scale Discovery plate imager and Daniela Farkas, BSc for her technical advice and assistance running our samples.

## Sources of Support

This work was funded by The Ohio State University College of Medicine Office of Research Dean’s Discovery Program (MNB and JAE). This work was also supported by funding from the National Heart, Lung, and Blood Institute (NHLBI) grants R01 HL155095 and R01 HL158532 (RDB) and National Cancer Institute grant P30 CA016058 (Genomics Shared Resource, OSUCCC). The results herein are based upon data generated by the LungMAP Consortium and downloaded from (www.lungmap.net) on May 3, 2023. The LungMAP consortium and the LungMAP Data Coordinating Center (U24-HL148865) are funded by NHLBI.

## Disclosures

The authors declare no conflicts of interest.

## Data availability statement

The raw sequencing data has been uploaded to the National Center for Biotechnology Information Gene Expression Omnibus database (NCBI GEO), accession number [data deposition pending]. All other data will be made available upon request.

## Notes

### Competing Interest Statement

The authors have declared no competing interest.

## References Cited

1. Forum of International Respiratory Societies. The Global Impact of Respiratory Disease - Third Edition. European Respiratory Society; 2021. at <firsnet.org/images/publications/FIRS_Master_09202021.pdf>.

2. Zhang J (Jim), Wei Y, Fang Z. Ozone Pollution: A Major Health Hazard Worldwide. Front Immunol 2019;10:2518.

3. Anenberg SC, Henze DK, Tinney V, Kinney PL, Raich W, Fann N, et al. Estimates of the Global Burden of Ambient PM2.5, Ozone, and NO2 on Asthma Incidence and Emergency Room Visits. Environ Health Perspect 2018;126:107004.

4. Khatri SB, Holguin FC, Ryan PB, Mannino D, Erzurum SC, Teague WG. Association of Ambient Ozone Exposure with Airway Inflammation and Allergy in Adults with Asthma. J Asthma 2009;46:777–785.

5. Camoretti-Mercado B, Lockey RF. Airway smooth muscle pathophysiology in asthma. Journal of Allergy and Clinical Immunology 2021;147:1983–1995.

6. Peters EA, Hiltermann JTN, Stolk J. Effect of apocynin on ozone-induced airway hyperresponsiveness to methacholine in asthmatics. Free Radical Biology and Medicine 2001;31:1442–1447.

7. Arjomandi M, Witten A, Abbritti E, Reintjes K, Schmidlin I, Zhai W, et al. Repeated Exposure to Ozone Increases Alveolar Macrophage Recruitment into Asthmatic Airways. Am J Respir Crit Care Med 2005;172:427–432.

8. Flayer CH, Ge MQ, Hwang JW, Kokalari B, Redai IG, Jiang Z, et al. Ozone Inhalation Attenuated the Effects of Budesonide on Aspergillus fumigatus-Induced Airway Inflammation and Hyperreactivity in Mice. Front Immunol 2019;10:.

9. Milara J, Navarro A, Almudéver P, Lluch J, Morcillo EJ, Cortijo J. Oxidative stress-induced glucocorticoid resistance is prevented by dual PDE3/PDE4 inhibition in human alveolar macrophages. Clinical & Experimental Allergy 2011;41:535–546.

10. Henriquez AR, Snow SJ, Schladweiler MC, Miller CN, Dye JA, Ledbetter AD, et al. Exacerbation of ozone-induced pulmonary and systemic effects by β2-adrenergic and/or glucocorticoid receptor agonist/s. Sci Rep 2019;9:17925.

11. Ho K, Weimar D, Torres-Matias G, Lee H, Shamsi S, Shalosky E, et al. Ozone impairs endogenous compensatory responses in allergic asthma. Toxicology and Applied Pharmacology 2023;459:116341.

12. Hatch GE, McKee J, Brown J, McDonnell W, Seal E, Soukup J, et al. Biomarkers of Dose and Effect of Inhaled Ozone in Resting versus Exercising Human Subjects: Comparison with Resting Rats. Biomark□Insights 2013;8:BMI.S11102.

13. Howenstine H, Eigen H, Tepper R. Pulmonary function in infants after pertussis. The Journal of Pediatrics 1991;118:563–566.

14. Ballinger MN, Paine R, Serezani CHC, Aronoff DM, Choi ES, Standiford TJ, et al. Role of Granulocyte Macrophage Colony-Stimulating Factor during Gram-Negative Lung Infection with Pseudomonas aeruginosa. Am J Respir Cell Mol Biol 2006;34:766–774.

15. Satija R, Farrell JA, Gennert D, Schier AF, Regev A. Spatial reconstruction of single-cell gene expression data. Nat Biotechnol 2015;33:495–502.

16. Germain P-L, Lun A, Garcia Meixide C, Macnair W, Robinson MD. Doublet identification in single-cell sequencing data using scDblFinder. F1000Res 2022;10:979.

17. Korsunsky I, Millard N, Fan J, Slowikowski K, Zhang F, Wei K, et al. Fast, sensitive and accurate integration of single-cell data with Harmony. Nat Methods 2019;16:1289–1296.

18. 18. Butler A, Darby C, Hao Y, Hartman A, Hoffman P, Jain J, et al. Azimuth: A Shiny App Demonstrating a Query-Reference Mapping Algorithm for Single-Cell Data. https://github.com/satijalab/azimuth 2023;at <https://github.com/satijalab/azimuth>.

19. Gaddis N, Fortriede J, Guo M, Bardes EE, Kouril M, Tabar S, et al. LungMAP Portal Ecosystem: Systems-level Exploration of the Lung. Am J Respir Cell Mol Biol 2024;70:129– 139.

20. Hicks SC, Liu R, Ni Y, Purdom E, Risso D. mbkmeans: Fast clustering for single cell data using mini-batch k-means. PLoS Comput Biol 2021;17:e1008625.

21. Wickham H. ggplot2. Cham: Springer International Publishing; 2016. doi:10.1007/978-3-319-24277-4.

22. Huber W, Carey VJ, Gentleman R, Anders S, Carlson M, Carvalho BS, et al. Orchestrating high-throughput genomic analysis with Bioconductor. Nat Methods 2015;12:115–121.

23. Robinson MD, McCarthy DJ, Smyth GK. edgeR: a Bioconductor package for differential expression analysis of digital gene expression data. Bioinformatics 2010;26:139–140.

24. Patial S, Saini Y. Lung macrophages: current understanding of their roles in Ozone-induced lung diseases. Crit Rev Toxicol 2020;50:310–323.

25. Jakubzick C, Gautier EL, Gibbings SL, Sojka DK, Schlitzer A, Johnson TE, et al. Minimal Differentiation of Classical Monocytes as They Survey Steady-State Tissues and Transport Antigen to Lymph Nodes. Immunity 2013;39:599–610.

26. Liu C-L, Shi G-P. Calcium-activated chloride channel regulator 1 (CLCA1): More than a regulator of chloride transport and mucus production. World Allergy Organization Journal 2019;12:100077.

27. Ito T, Hirose K, Saku A, Kono K, Takatori H, Tamachi T, et al. IL-22 induces Reg3γ and inhibits allergic inflammation in house dust mite–induced asthma models. J Exp Med 2017;214:3037–3050.

28. Dodd KM, Yang J, Shen MH, Sampson JR, Tee AR. mTORC1 drives HIF-1α and VEGF-A signalling via multiple mechanisms involving 4E-BP1, S6K1 and STAT3. Oncogene 2015;34:2239–2250.

29. Zoungrana LI, Krause-Hauch M, Wang H, Fatmi MK, Bates L, Li Z, et al. The Interaction of mTOR and Nrf2 in Neurogenesis and Its Implication in Neurodegenerative Diseases. Cells 2022;11:2048.

30. Stevens NC, Brown VJ, Domanico MC, Edwards PC, Van Winkle LS, Fiehn O. Alteration of glycosphingolipid metabolism by ozone is associated with exacerbation of allergic asthma characteristics in mice. Toxicological Sciences 2023;191:79–89.

31. Maguire TJA, Yung S, Ortiz-Zapater E, Kayode OS, Till S, Corrigan C, et al. Sphingosine-1-phosphate induces airway smooth muscle hyperresponsiveness and proliferation. Journal of Allergy and Clinical Immunology 2023;152:1131–1140.e6.

32. Chung F-T, Huang H-Y, Lo C-Y, Huang Y-C, Lin C-W, He C-C, et al. Increased Ratio of Matrix Metalloproteinase-9 (MMP-9)/Tissue Inhibitor Metalloproteinase-1 from Alveolar Macrophages in Chronic Asthma with a Fast Decline in FEV1 at 5-Year Follow-up. Journal of Clinical Medicine 2019;8:1451.

33. Misharin AV, Morales-Nebreda L, Reyfman PA, Cuda CM, Walter JM, McQuattie-Pimentel AC, et al. Monocyte-derived alveolar macrophages drive lung fibrosis and persist in the lung over the life span. Journal of Experimental Medicine 2017;jem.20162152. doi:10.1084/jem.20162152.

34. Zasłona Z, Przybranowski S, Wilke C, Rooijen N van, Teitz-Tennenbaum S, Osterholzer JJ, et al. Resident Alveolar Macrophages Suppress, whereas Recruited Monocytes Promote, Allergic Lung Inflammation in Murine Models of Asthma. The Journal of Immunology 2014;193:4245–4253.

35. Guttenberg MA, Vose AT, Birukova A, Lewars K, Cumming RI, Albright MC, et al. Tissue-Resident Alveolar Macrophages Reduce Ozone-induced Inflammation via MerTK Mediated Efferocytosis. Am J Respir Cell Mol Biol 2024;doi:10.1165/rcmb.2023-0390OC.

36. Moon H-G, Kim S, Kim K-H, Kim Y-M, Rehman J, Lee H, et al. CX3CR1+ Macrophage Facilitates the Resolution of Allergic Lung Inflammation via Interacting CCL26. Am J Respir Crit Care Med 2023;207:1451–1463.

37. Wohnhaas CT, Baßler K, Watson CK, Shen Y, Leparc GG, Tilp C, et al. Monocyte-derived alveolar macrophages are key drivers of smoke-induced lung inflammation and tissue remodeling. Front Immunol 2024;15:.

38. Bagato O, Balkema-Buschmann A, Todt D, Weber S, Gömer A, Qu B, et al. Spatiotemporal analysis of SARS-CoV-2 infection reveals an expansive wave of monocyte-derived macrophages associated with vascular damage and virus clearance in hamster lungs. Microbiology Spectrum 2023;12:e02469–23.

39. Liao M, Liu Y, Yuan J, Wen Y, Xu G, Zhao J, et al. Single-cell landscape of bronchoalveolar immune cells in patients with COVID-19. Nat Med 2020;26:842–844.

40. Merad M, Martin JC. Pathological inflammation in patients with COVID-19: a key role for monocytes and macrophages. Nat Rev Immunol 2020;20:355–362.

41. Hou F, Wang H, Zheng K, Yang W, Xiao K, Rong Z, et al. Distinct Transcriptional and Functional Differences of Lung Resident and Monocyte-Derived Alveolar Macrophages During the Recovery Period of Acute Lung Injury. Immune Network 2023;23:.

42. Lee YG, Jeong JJ, Nyenhuis S, Berdyshev E, Chung S, Ranjan R, et al. Recruited Alveolar Macrophages, in Response to Airway Epithelial–Derived Monocyte Chemoattractant Protein 1/CCL2, Regulate Airway Inflammation and Remodeling in Allergic Asthma. American Journal of Respiratory Cell and Molecular Biology 2015;52:772–784.

43. Mellado M, Ana AM de, Gómez L, Martínez-A C, Rodríguez-Frade JM. Chemokine Receptor 2 Blockade Prevents Asthma in a Cynomolgus Monkey Model. J Pharmacol Exp Ther 2008;324:769–775.

44. Enweasor C, Flayer CH, Haczku A. Ozone-Induced Oxidative Stress, Neutrophilic Airway Inflammation, and Glucocorticoid Resistance in Asthma. Frontiers in Immunology 2021;12:.

45. O’Byrne P m., Walters E h., Gold B d., Aizawa H a., Fabbri L m., Alpert S e., et al. Neutrophil Depletion Inhibits Airway Hyperresponsiveness Induced by Ozone Exposure. Am Rev Respir Dis 1984;130:214–219.

46. Matsumoto K, Aizawa H, Inoue H, Koto H, Nakano H, Hara N. Role of neutrophil elastase in ozone-induced airway responses in guinea-pigs. European Respiratory Journal 1999;14:1088–1094.

47. Koga H, Miyahara N, Fuchimoto Y, Ikeda G, Waseda K, Ono K, et al. Inhibition of neutrophil elastase attenuates airway hyperresponsiveness and inflammation in a mouse model of secondary allergen challenge: neutrophil elastase inhibition attenuates allergic airway responses. Respir Res 2013;14:8.

48. Hiltermann TJN, Peters EA, Alberts B, Kwikkers K, Borggreven PA, Hiemstra PS, et al. Ozone-Induced Airway Hyperresponsiveness in Patients With Asthma: Role of Neutrophil-Derived Serine Proteinases. Free Radical Biology and Medicine 1998;24:952–958.

49. Taube C, Nick JA, Siegmund B, Duez C, Takeda K, Rha Y-H, et al. Inhibition of Early Airway Neutrophilia Does Not Affect Development of Airway Hyperresponsiveness. Am J Respir Cell Mol Biol 2004;30:837–843.

50. Kawano H, Kayama H, Nakama T, Hashimoto T, Umemoto E, Takeda K. IL-10-producing lung interstitial macrophages prevent neutrophilic asthma. Int Immunol 2016;28:489–501.

51. Tighe RM, Birukova A, Malakhau Y, Kobayashi Y, Vose AT, Chandramohan V, et al. Altered Ontogeny and Transcriptomic Signatures of Tissue-Resident Pulmonary Interstitial Macrophage Ameliorate Allergic Airway Hyperresponsiveness. Front Immunol 2024;15:.

52. Schmidt D, Rabe KF. Immune mechanisms of smooth muscle hyperreactivity in asthma. Journal of Allergy and Clinical Immunology 2000;105:673–682.

53. Britt RD, Ruwanpathirana A, Ford ML, Lewis BW. Macrophages Orchestrate Airway Inflammation, Remodeling, and Resolution in Asthma. Int J Mol Sci 2023;24:10451.

54. Manson ML, Säfholm J, James A, Johnsson A-K, Bergman P, Al-Ameri M, et al. IL-13 and IL-4, but not IL-5 nor IL-17A, induce hyperresponsiveness in isolated human small airways. J Allergy Clin Immunol 2020;145:808–817.e2.

55. Evans CM, Fryer AD, Jacoby DB, Gleich GJ, Costello RW. Pretreatment with antibody to eosinophil major basic protein prevents hyperresponsiveness by protecting neuronal M2 muscarinic receptors in antigen-challenged guinea pigs. J Clin Invest 1997;100:2254–2262.

56. Yost BL, Gleich GJ, Fryer AD. Ozone-induced hyperresponsiveness and blockade of M2 muscarinic receptors by eosinophil major basic protein. Journal of Applied Physiology 1999;87:1272–1278.

57. Fang X, Huang S, Zhu Y, Lei J, Xu Y, Niu Y, et al. Short-term exposure to ozone and asthma exacerbation in adults: A longitudinal study in China. Front Public Health 2023;10:1070231.

58. Most Recent National Asthma Data | CDC. 2024;at <https://www.cdc.gov/asthma/most_recent_national_asthma_data.htm>.

59. Downie SR, Salome CM, Verbanck S, Thompson B, Berend N, King GG. Ventilation heterogeneity is a major determinant of airway hyperresponsiveness in asthma, independent of airway inflammation. Thorax 2007;62:684–689.

60. Tliba O, Panettieri RA. Paucigranulocytic asthma: The uncoupling of airway obstruction from inflammation. The Journal of allergy and clinical immunology 2019;143:1287.

