## Supplemental Data for "scRNA-seq identifies unique macrophage population in murine model of ozone induced asthma exacerbation"

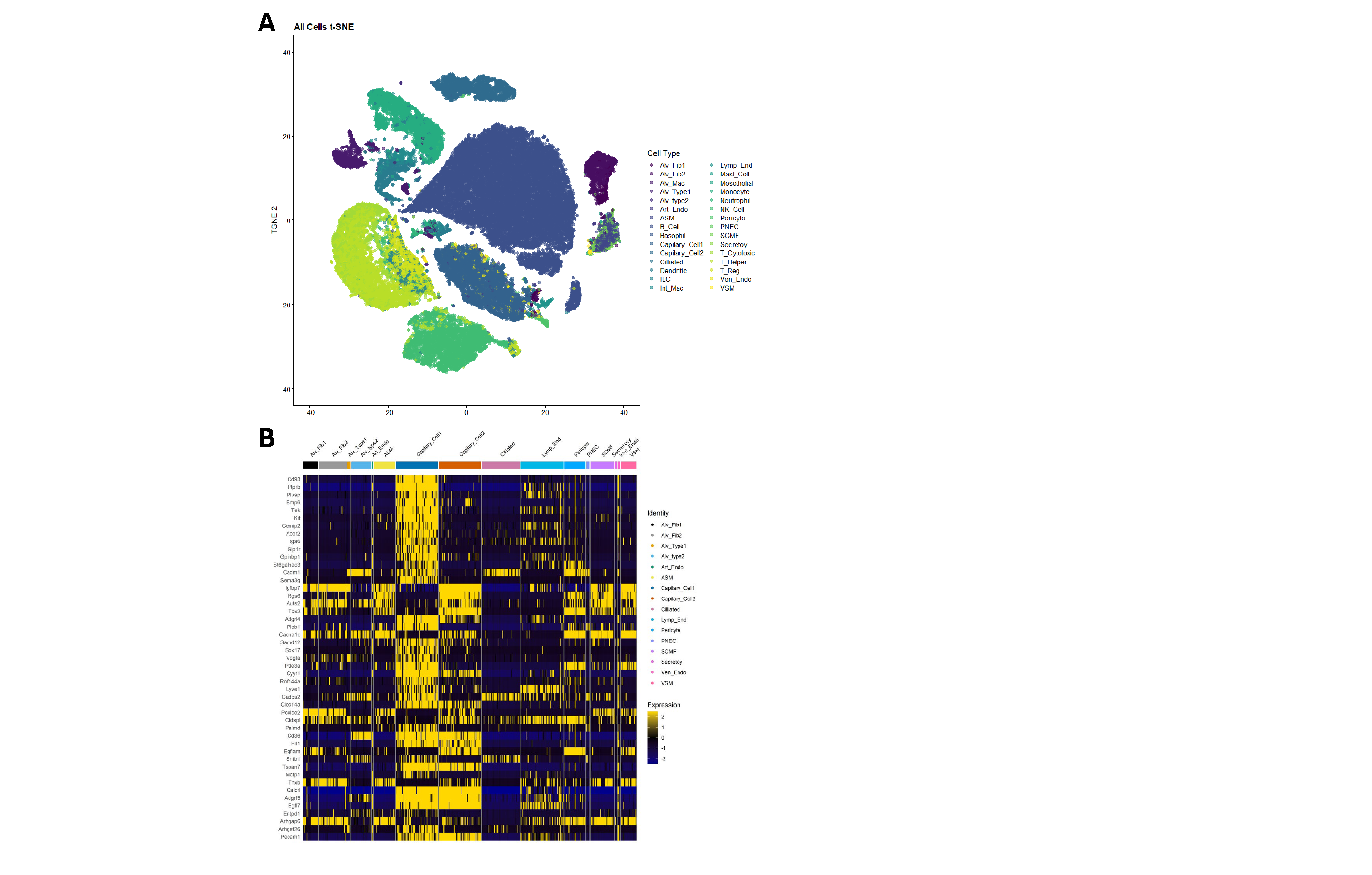
**Supplemental Figure 1.** *Identification of cell clusters from mouse lungs using single cell RNA sequencing.* Male and female C57BL/6 mice were exposed to DRA, O_3_, or DRA+O_3_ as shown in Figure 1A. Whole lung tissue was collected and then digested using collagenase followed by live cell enrichment using FACS. Live cells were then processed for scRNA sequencing. (A) t-SNE plot of cell clusters identified in all 4 treatment groups (control, DRA, O_3_, DRA+O_3_) and (B) heatmap of previously validated marker genes used to annotate clusters.

**Supplemental Table 1.** Number of cells from scRNA sequencing cluster analysis.

| **Cell Type** | **Control** | **DRA** | **O_3_** | **DRA+O_3_** | **Total** |
| --- | --- | --- | --- | --- | --- |
| Alv_Mac | 384 | 409 | 353 | 359 | **1,505** |
| B_Cell | 5,491 | 5,403 | 5,194 | 2,984 | **19,072** |
| Dendritic | 218 | 481 | 190 | 458 | **1,347** |
| Monocyte | 792 | 1,386 | 696 | 764 | **3,638** |
| NK_Cell | 1,113 | 1,355 | 1,385 | 708 | **4,561** |
| T_Cytotoxic | 386 | 666 | 721 | 394 | **2,167** |
| T_Helper | 884 | 1,549 | 1,135 | 698 | **4,266** |
| T_Reg | 80 | 515 | 92 | 601 | **1,288** |
| ILC | 95 | 206 | 139 | 123 | **563** |
| Int_Mac | 33 | 96 | 53 | 213 | **395** |
| Neutrophil | 40 | 62 | 21 | 91 | **214** |
| Basophil | 12 | 10 | 1 | 9 | **32** |
| Mast_Cell | 0 | 0 | 2 | 0 | **2** |
| Capilary_Cell1 | 1,450 | 918 | 1,209 | 1,398 | **4,975** |
| Capilary_Cell2 | 653 | 446 | 604 | 522 | **2,225** |
| Alv_Fib2 | 492 | 167 | 395 | 107 | **1,161** |
| SCMF | 345 | 84 | 125 | 82 | **636** |
| ASM | 110 | 35 | 231 | 96 | **472** |
| Alv_Fib1 | 176 | 58 | 71 | 98 | **403** |
| Ven_Endo | 91 | 20 | 56 | 65 | **232** |
| Pericyte | 91 | 41 | 26 | 55 | **213** |
| Lymp_End | 71 | 27 | 55 | 32 | **185** |
| Art_Endo | 33 | 6 | 15 | 25 | **79** |
| VSM | 30 | 9 | 27 | 6 | **72** |
| Ciliated | 17 | 11 | 10 | 8 | **46** |
| Alv_type2 | 1 | 9 | 6 | 8 | **24** |
| Mesothelial | 1 | 6 | 10 | 5 | **22** |
| PNEC | 2 | 0 | 2 | 1 | **5** |
| Alv_Type1 | 2 | 1 | 0 | 1 | **4** |
| Secretory | 1 | 1 | 0 | 0 | **2** |

**Supplemental Table 2.** Number of DEGs in immune cells.

| **Cell Type** | **DRA vs Control** | | | **O_3_ vs Control** | | | **DRA+O_3_ vs Control** | | |
| --- | --- | --- | --- | --- | --- | --- | --- | --- | --- |
|  | **up** | **down** | **total** | **up** | **down** | **total** | **up** | **down** | **total** |
| Alv_Mac | 298 | 128 | 426 | 90 | 55 | 145 | 723 | 624 | 1347 |
| B_Cell | 15 | 1 | 16 | 31 | 64 | 95 | 64 | 8 | 72 |
| Dendritic | 124 | 122 | 246 | 98 | 68 | 166 | 288 | 291 | 579 |
| Monocyte | 24 | 11 | 35 | 38 | 141 | 179 | 74 | 17 | 91 |
| NK_Cell | 11 | 5 | 16 | 15 | 10 | 25 | 34 | 6 | 40 |
| T_Cytotoxic | 42 | 3 | 45 | 25 | 16 | 41 | 139 | 22 | 161 |
| T_Helper | 35 | 10 | 45 | 50 | 17 | 67 | 72 | 7 | 79 |
| T_Reg | 140 | 50 | 190 | 11 | 27 | 38 | 235 | 87 | 322 |

Differentially expressed genes (DEGs) for immune cell clusters with over 1,000 total cells. DEG defined as FDR < .05 and log2(fold change) ≥ |1.0|.

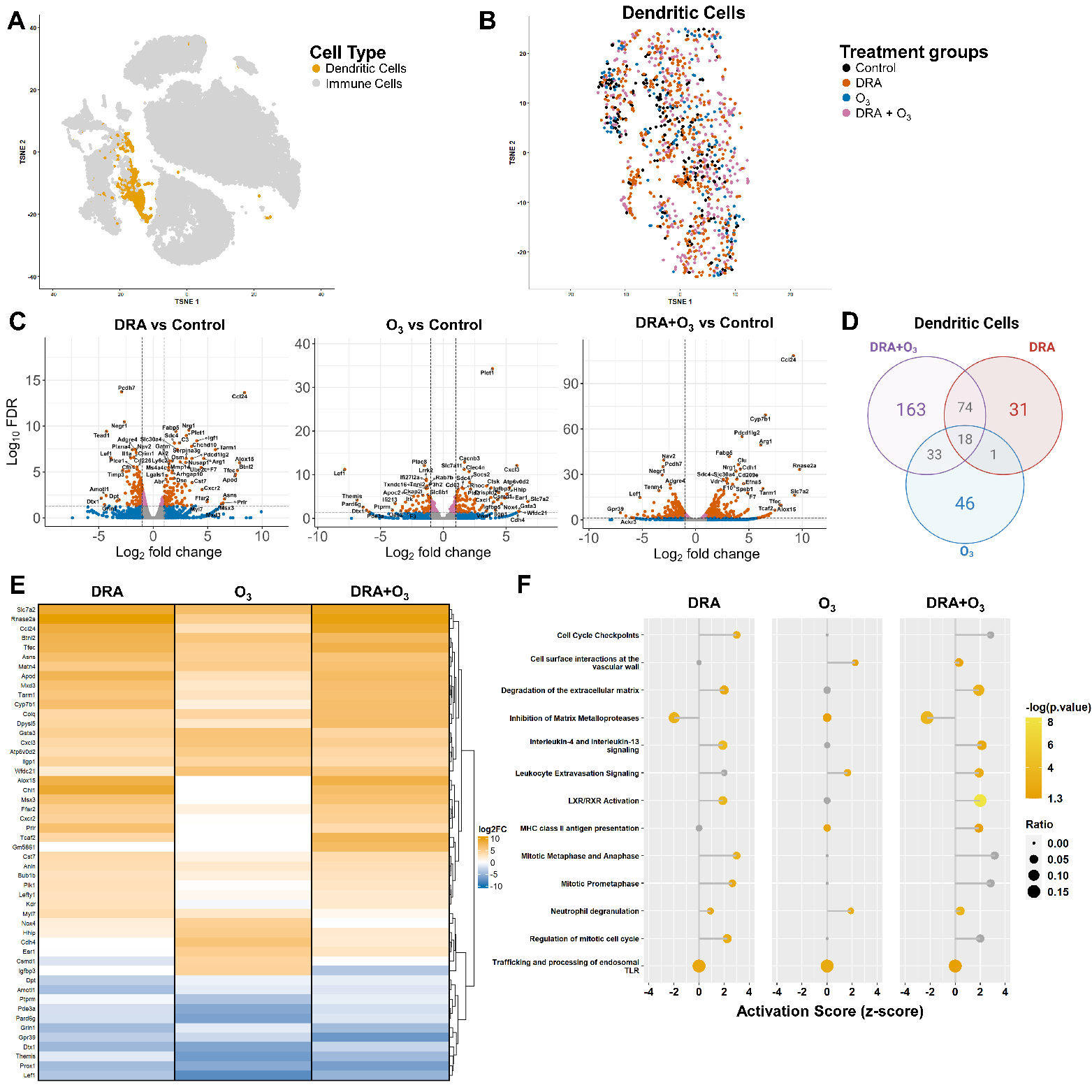
**Supplemental Figure 2.** *scRNA sequencing of DCs.* (A) t-SNE plot of all immune cell clusters identified in the lungs of control, DRA, O_3_, and DRA+O_3_ mice, with DCs highlighted in orange. (B) t-SNE plot of DCs from all four groups; colors denote exposure(s) as indicated. (C) Volcano plots of differentially expressed genes (DEG) in DCs from DRA, O_3_, or DRA+O_3_ exposed mice (respectively) compared to DCs from the control group. Each dot represents one gene; “significant” defined as Log_10_(false discover rate [FDR]) < .05 and “relevant” mean expression level defined as Log_2_(fold change) > |1.0|, genes meeting both criteria considered differentially expressed (“relevant” red dots above horizontal significance dashed line and outside vertical relevance dashed lines). (D) Venn diagram of the number of unique and overlapping DEG (upregulated only) in DCs from each treatment group compared to controls. (E) Heatmap displaying the top DEG in DCs from respective treatment groups compared to controls. (F) Dot plot displaying transcriptional pathway enrichment analysis in DCs from respective treatment groups compared to control DCs. The x-axis denotes respective pathway activation scores (z-score ≤ -2 considered inhibited and ≥ 2 considered activated) and dot color corresponds to -log(p-value); -log(0.05) = 1.3, therefore pathways with -log(p-value) ≤ 1.3 are considered non-significant, as indicated by grey dot color. The dot size is proportionate to DEG overlap ratio [DEG in data set/total genes in pathway].

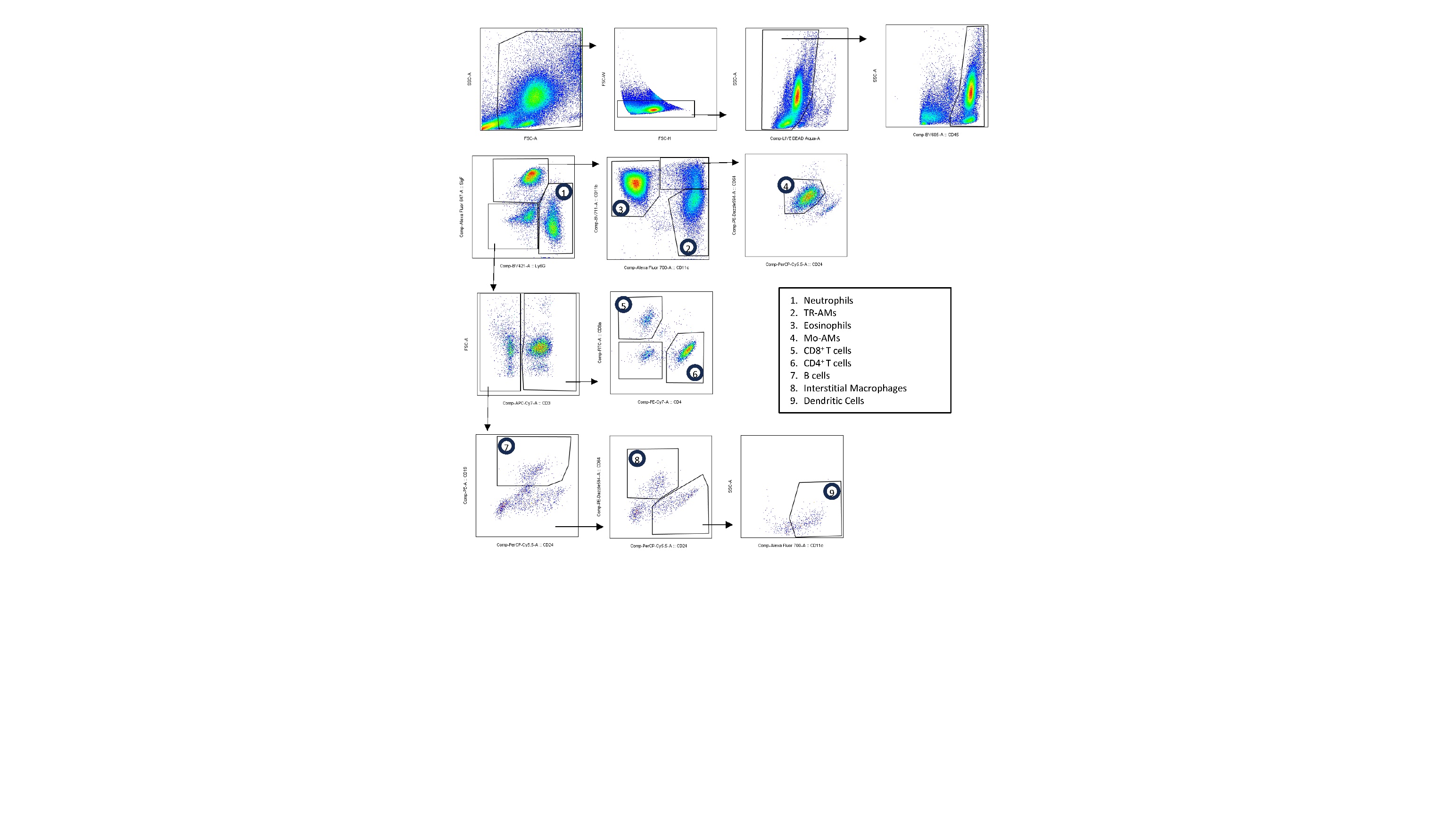
**Supplemental Figure 3.** *Flow Cytometry gating scheme.* Representative gating for the identification of immune cell populations in the airspaces of control, DRA, O_3_, and DRA+O_3_ treated mice (Figure 1A). Cells were first gated for size, doublet exclusion, viability, CD45^+^ and further defined as follows: neutrophils (Siglec F^-^ Ly6G^+^ CD11b^+^), eosinophils (Siglec F^+^ CD11b^+^ CD11c^-^ CD24^+^), TR-AMs (Siglec F^+^ CD11b^-^ CD11c^+^ CD24^-^ CD64^+^), Mo-AMs (Siglec F^mid/low^ CD11b^+^ CD11c^+^ CD24^-^ CD64^+^), interstitial macrophages (Siglec F^-^ CD64^+^), dendritic cells (Siglec F^-^ CD11c^+^ CD24^+^), T cells (Siglec F^-^ CD3^+^ CD4^+/-^ CD8^+/-^), and B cells (Siglec F^-^ CD3^-^ CD19^+^ CD24^+^). The total number of each cell type was calculated by multiplying the population frequency (% of all BAL cells) by the total number of BAL cells collected.

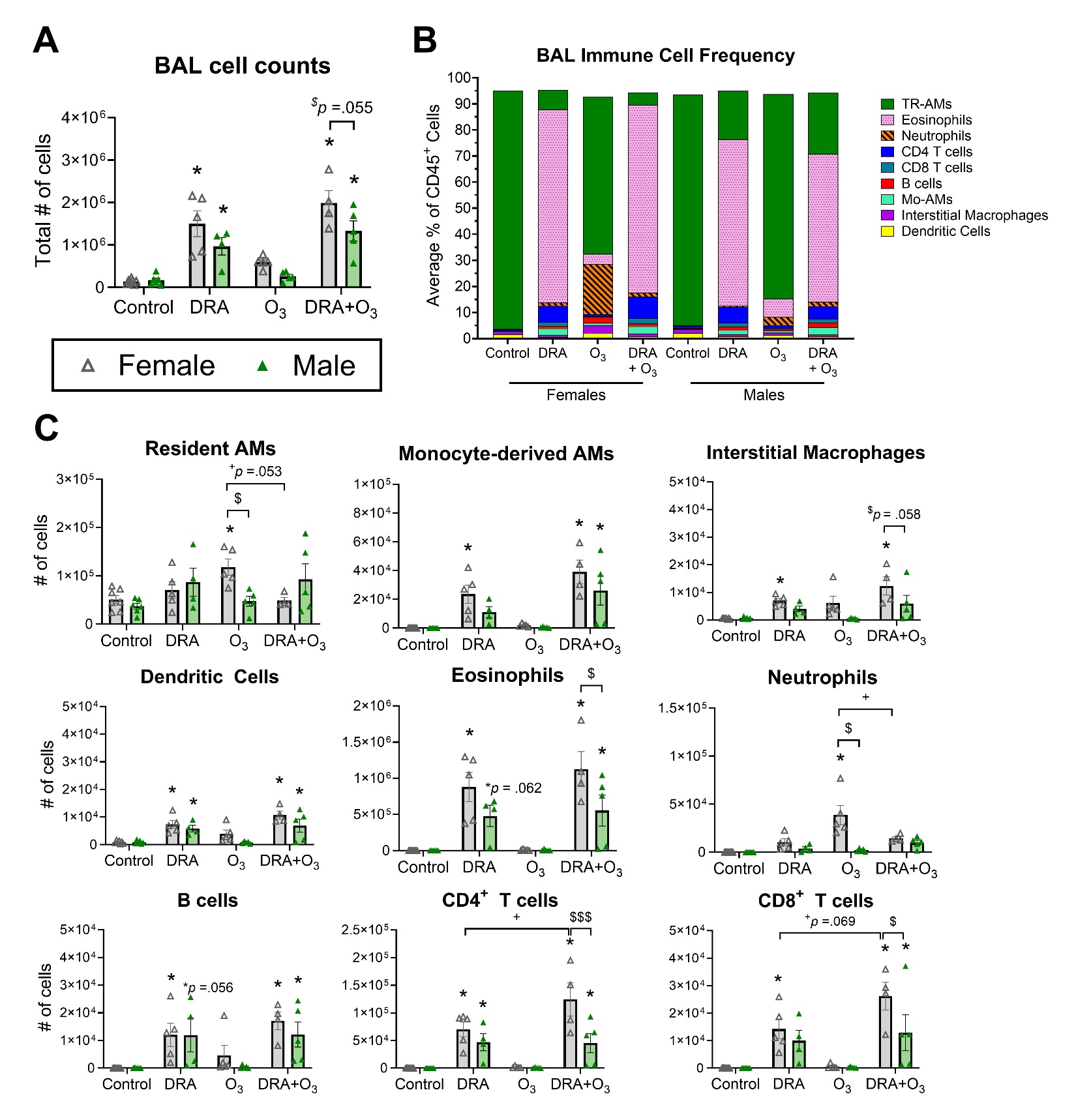

**Supplemental Figure 4**. *Sex-based comparison of airspace immune cell populations.* Sex-based analysis of cells within the airspaces collected for (A) total cell counts, and flow cytometry used to determine (B) frequency and (C) total number of immune cell population in the BAL. *n* = 4-5; **p* < .05 compared to same-sex controls, ^$^*p* < .05 compared to the opposite sex of the same treatment, ^+^*p* < .05 compared to the DRA+O_3_ group of the same sex. Statistical significance analyzed using two-way ANOVA followed by Tukey’s and Sidak’s *post hoc* testing. All data presented as *mean ± SEM*. BAL, bronchoalveolar lavage.

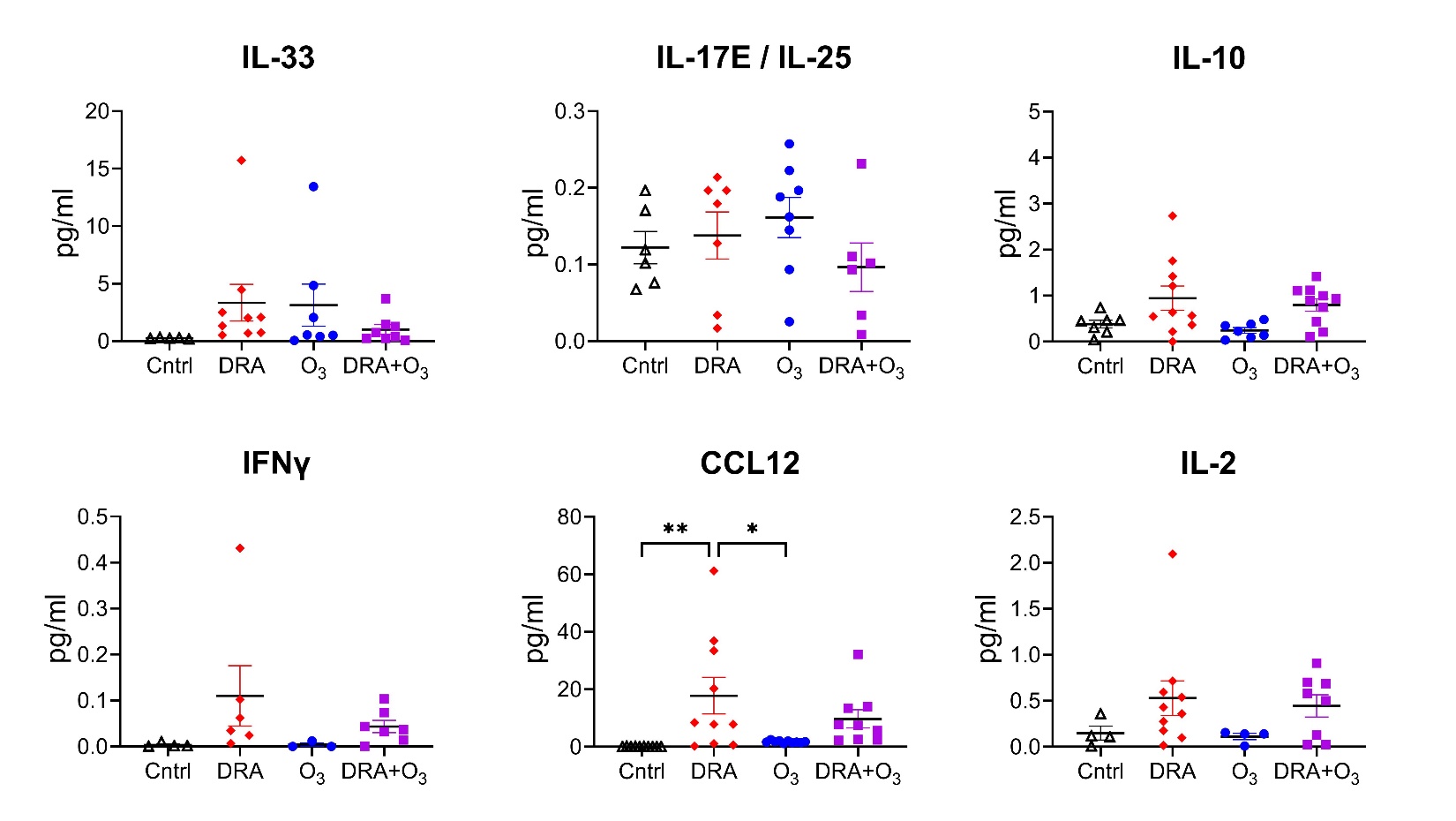

**Supplemental Figure 5.** *Additional airspace cytokines in DRA, O_3_, and DRA+O_3_ exposed mice.* As shown in Figure 1A, male and female mice were intranasally sensitized and challenged with triple allergen mixture (DRA) five times over 17 days; 24 hr after the last DRA administration, mice were exposed to 2 ppm O_3_ for 3 hr; 24 hr after O_3_ concentrated bronchoalveolar lavage fluid was collected for airspace cytokine measurements. All data presented as *mean ± SEM*; *n* = 9-10; **p* < .05, ***p* < .01, ****p* < .001 as indicated; statistical significance analyzed using one-way ANOVA followed by Tukey’s *post hoc* testing.

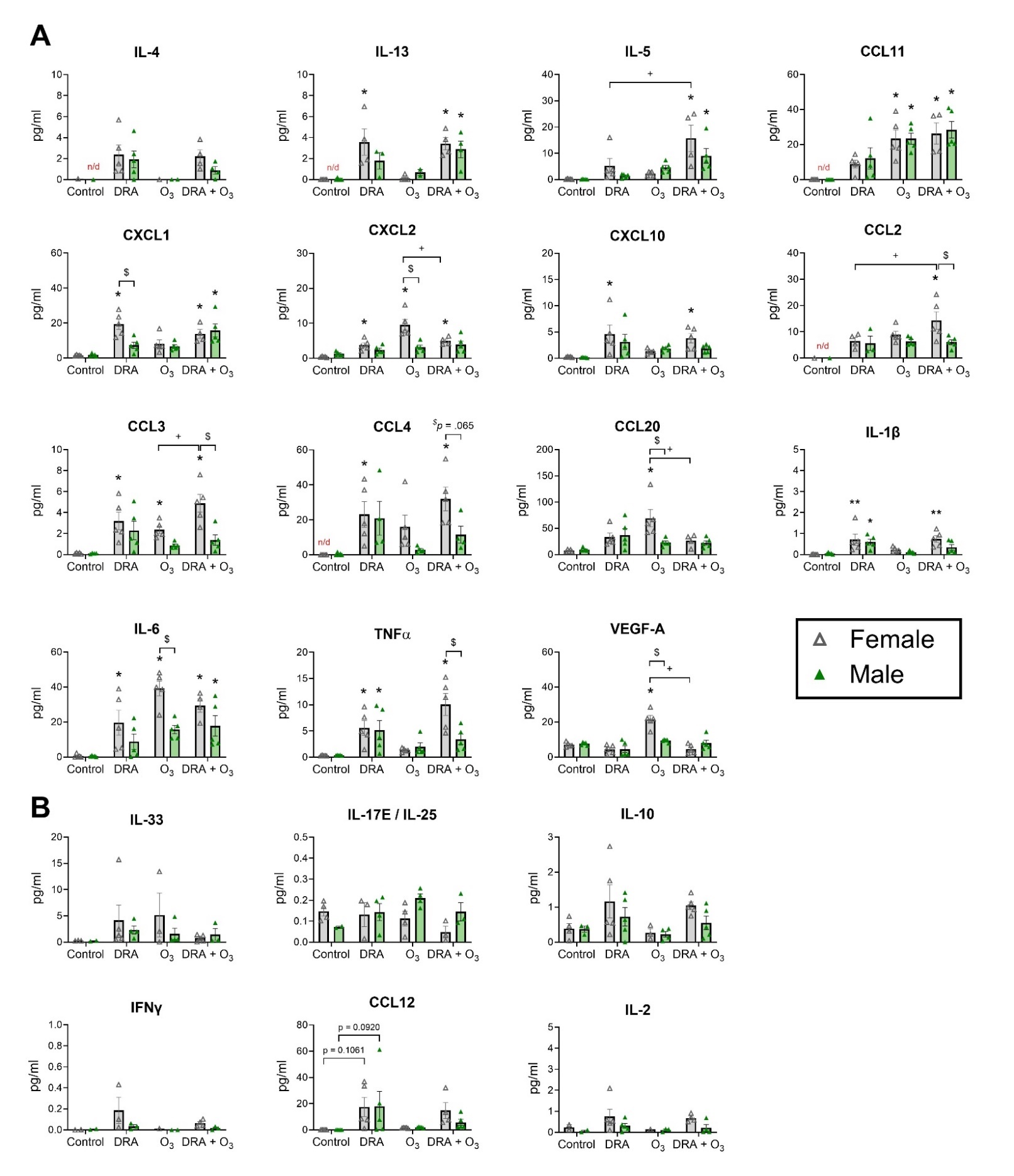
**Supplemental Figure 6**. *Sex-based comparison of airspace cytokines.* Sex-based analysis of (A) cytokines listed in main text Figure 5, and (B) cytokines found in supplemental figure 5. All data presented as *mean ± SEM*; *n* = 4-5; **p* < .05 compared to same-sex controls, ^$^*p* < .05 compared to the opposite sex of the same treatment, ^+^*p* < .05 compared to the DRA+O_3_ group of the same sex. Statistical significance analyzed using two-way ANOVA followed by Tukey’s and Sidak’s *post hoc* testing.

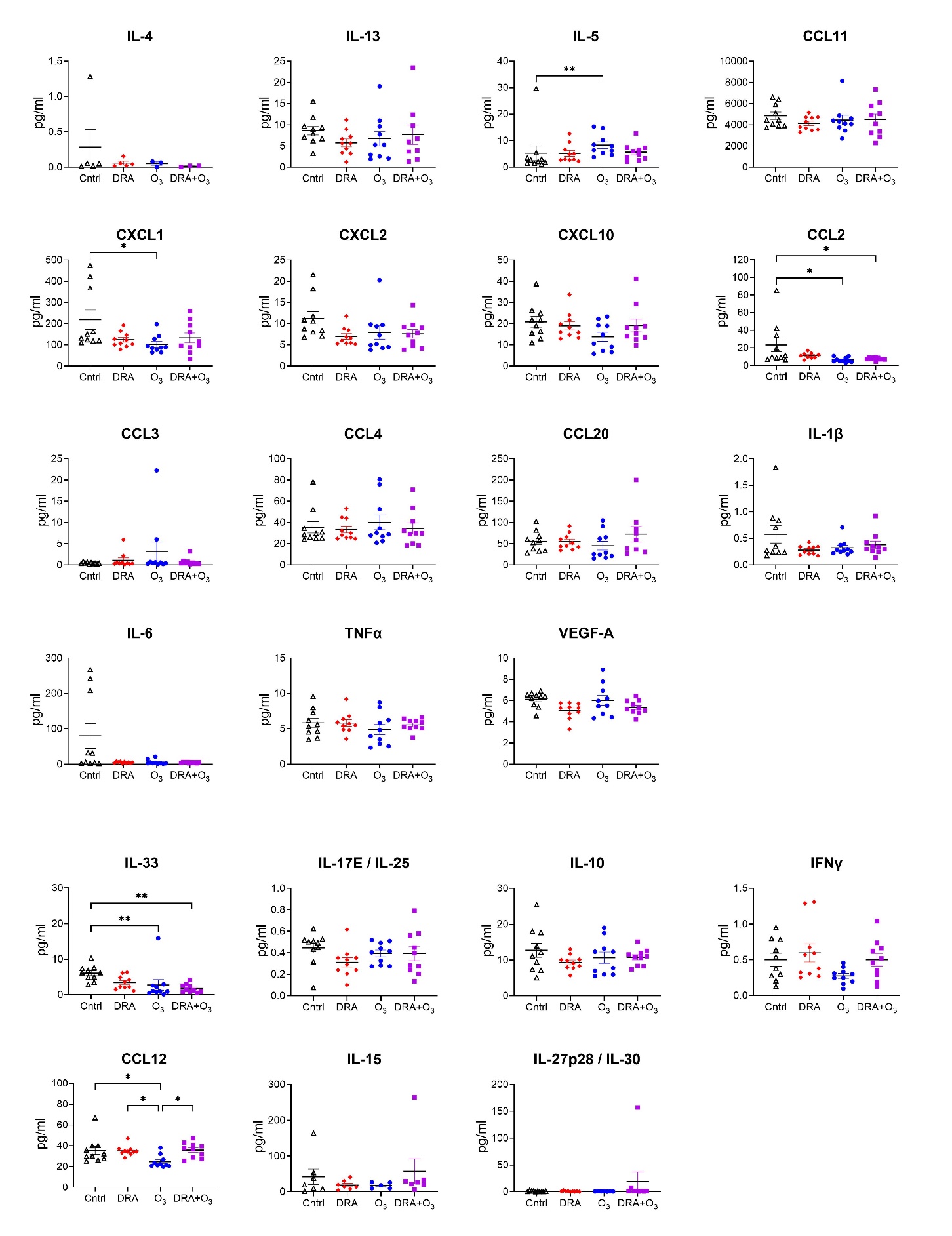
**Supplemental Figure 7.** *Plasma cytokine levels in DRA, O_3_, and DRA+O_3_ exposed mice.* Mice were exposed as shown in Figure 1A, and heparinized blood was collected for plasma cytokine measurements. All data presented as *mean ± SEM*; *n* = 9-10; **p* < .05, ***p* < .01 as indicated; statistical significance analyzed using one-way ANOVA followed by Tukey’s *post hoc* testing.
